# A Computational Framework for Understanding HCV Infection and Associated Lipid Dynamics

**DOI:** 10.64898/2026.10.16.759547

**Authors:** Andrew Jacoby, Samiran Ghosh, Malay Banerjee, Aditi Ghosh, Vitaly Volpert

## Abstract

Hepatitis C virus (HCV) infection is strongly intertwined with host lipid metabolism and is frequently associated with hepatic steatosis, particularly in genotype-specific cases where lipid accumulation reflects direct viral effects rather than host metabolic comorbidities. Despite extensive clinical and experimental evidence, a quantitative framework capturing the bidirectional coupling between viral dynamics and lipid metabolism remains lacking. In this work, we develop a comprehensive mathematical model that integrates HCV infection dynamics with intracellular lipid regulation. The framework combines a reaction diffusion system for spatial viral spread with a coupled metabolic module describing lipid influx. Numerical simulations highlight the role of lipid feedback in shifting infection outcomes toward persistence and recurrent oscillations, demonstrating that lipid accumulation acts as a critical driver of viral amplification. The model also highlights the critical role of immune memory to study acute and chronic phase and successfully reproduces observed viral RNA trajectories. Overall, the framework provides mechanistic insights into HCV dynamics and offers a general modeling approach for studying virus metabolism interactions in liver disease.

## 1 Introduction

Hepatitis C virus (HCV) remains a major cause of chronic liver disease worldwide and is tightly linked with host lipid metabolism throughout its life cycle. Clinically, hepatic steatosis is detected in 40–70 % of patients with chronic hepatitis C and is particularly frequent and severe in infections with genotype 3, where steatosis correlates with viral load and likely reflects direct cytopathic effects of the virus rather than host metabolic comorbidities [3, 10, 11, 14]. Beyond its histological footprint, steatosis is associated with insulin resistance and adverse disease progression, including accelerated fibrosis and higher risk of hepatocellular carcinoma, underlining the need for mechanistic understanding and quantitative frameworks that can disentangle virus host metabolic crosstalk [1, 6, 10, 11].

At the cellular level, HCV exploits and perturbs lipid pathways at multiple nodes. Viral entry depends on lipoprotein receptors and lipid components, replication and assembly are organized on endoplasmic reticulum (ER) membranes in close apposition to lipid droplets (LDs), and two key viral proteins core and NS5A (nonstructural protein 5A) localize to LDs to coordinate assembly or exit [1, 6, 9, 18]. HCV reprograms hepatocyte lipid homeostasis by activating sterol regulatory element binding proteins (SREBPs), suppressing peroxisome proliferator–activated receptor *α* (PPAR-*α*), and inhibiting microsomal triglyceride transfer protein (MTP). Thereby promoting new lipogenesis, diminishing *β*-oxidation, impairing very low-density lipoprotein (VLDL) assembly/secretion, and driving triglyceride sequestration into LDs [2,13,14,16,18] . Enzymes such as diacylglycerol acyltransferase-1 (DGAT1) further couple triglyceride synthesis to viral assembly by recruiting NS5A to LDs, linking lipid fluxes directly to infectious particle production [10]. Quantitative imaging studies indicate that HCV-associated lipid accumulation often reflects an increased number of LDs rather than larger droplet size, offering measurable phenotypes to benchmark models [16].

Mathematical modeling has been instrumental in describing HCV kinetics under therapy and the intracellular processes of replication and assembly. Seminal within host models explains biphasic viral load declines and informed treatment strategies by quantifying infection rates, clearance, and drug effects [15, 17, 19]. Intracellular models have since incorporated replication complexes and virion production, providing a scaffold for coupling metabolism to viral output [17]. In parallel, a rich systems-biology literature has emerged for hepatic lipid metabolism and steatosis including mechanistic ODE models of fatty acid handling, triglyceride synthesis, LD turnover, and VLDL assembly; spatially informed sinusoidal (zonated) models; and quantitative systems pharmacology (QSP) platforms for NAFLD/MASLD [4,7,15,17,19]. These models capture key control points such as MTP and Apo*β* dependent VLDL biogenesis, DGAT-mediated triglyceride formation, and the competition between storage (LDs), oxidation, and secretion precisely the pathways perturbed by HCV [7, 12]. Notably, mathematical treatments of MTP inhibition reproduce steatogenic phenotypes and provide a blueprint for embedding HCV-specific perturbations into lipid-flux networks [5].

Despite these advances, an integrated quantitative framework that jointly describes HCV dynamics and the host lipid economy underlying steatosis remains underdeveloped. Such a framework would enable (i) mechanistic attribution of steatosis to viral versus host metabolic drivers (with genotype-specific parameters), (ii) prediction of how antiviral or metabolic interventions propagate through lipid fluxes and viral outputs, and (iii) identification of measurable biomarkers e.g., LD number/turnover, VLDL-TG secretion that best constrain model parameters. In this work, we develop a virus metabolism coupling that links standard HCV infection dynamics to a hepatocyte lipid module.

We study here specifically how does the bidirectional interaction between HCV viral replication and host intracellular lipid metabolism mechanistically drive the onset, persistence, and spatial spread of viral steatosis. We also study what combination of viral, immune, and lipid-feedback parameters determines whether an infection resolves, becomes chronic, or oscillates persistently over time.

This article’s central contribution is the first mathematical framework that bidirectionally couples HCV viral dynamics with intracellular lipid metabolism. The model uniquely captures how HCV infection drives lipid accumulation via lipid influx while lipid accumulation simultaneously feeds back to amplify viral replication, formalized through a lipid-dependent viral production term. The framework extends this coupling into a spatial reaction-diffusion setting, yielding new analytical results for infection wave speed and spreading thresholds that incorporate immune clearance, and further shows numerically that long-term infection behavior is insensitive to initial spatial seeding. On the ODE side, the same model reproduces the full clinical spectrum of HCV outcomes spontaneous clearance, stable chronic infection, damped oscillatory infection, and sustained relapsing dynamics from a single parameter set varied along one axis *R*_0_, and identifies a previously uncharacterized nonzero threshold in immune memory depletion rate that governs the acute-to-chronic transition. The model is further validated by fitting to real clinical HCV RNA trajectories, grounding the theoretical predictions in observed patient data, and is proposed as a generalizable template for virus metabolism modeling in liver disease beyond HCV specifically.

The remainder of this paper is organized as follows. Section 2 presents the coupled HCV-lipid model and its biological foundations. Section 3 develops the ODE formulation and investigates equilibrium states, threshold conditions, stability, bifurcations, and the role of immune memory in acute and chronic infection dynamics. Section 4 analyzes the spatial PDE model, focusing on infection propagation, traveling-wave behavior, and numerical validation of infection spread. Finally, the paper concludes with a discussion of the biological implications and future research directions.

## 2 Model of spread of infection

HCV promotes viral steatosis due to (a) increase of lipid influx in the liver hepatocytes from the adipocyte tissue because virus initiates insulin resistance, (b) inhibition of beta-oxidation in hepatocytes, (c) inhibition of lipid transport. We consider the following model as in Figure 1:

**Figure 1:**
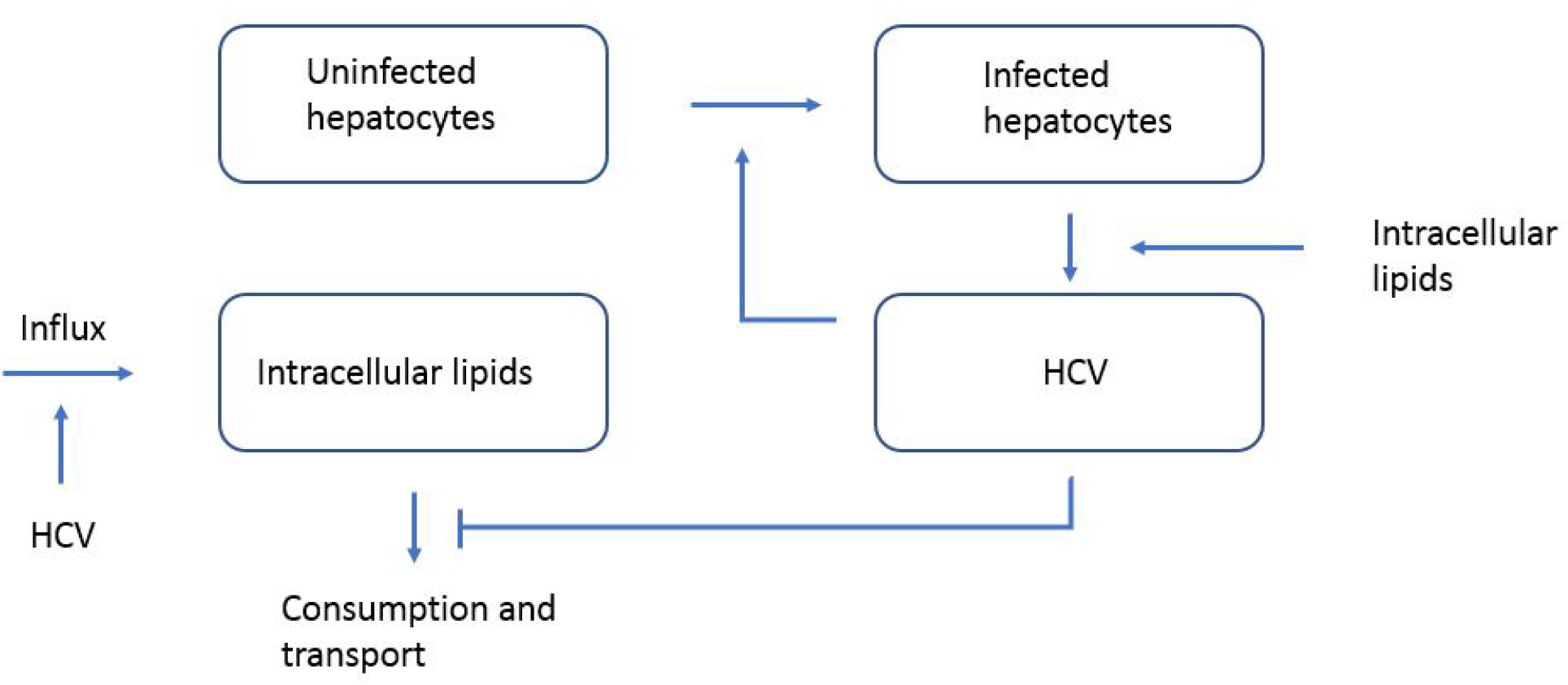
Schematic representation of the model. HCV infects uninfected cells. Infected cells produce virus. Virus up-regulates influx of lipids in the liver due to imposed insulin resistance, and down-regulates intracellular lipid consumption and transport, leading to its excessive accumulation and, as a consequence, steatosis and inflammation. On the other hand, lipid accumulation stimulates virus replication.

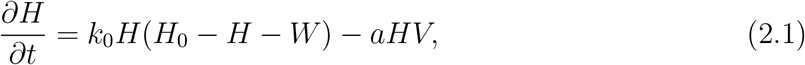

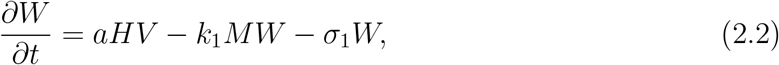

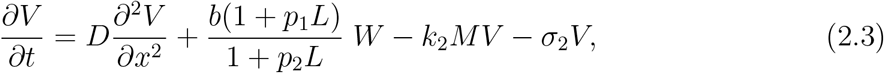

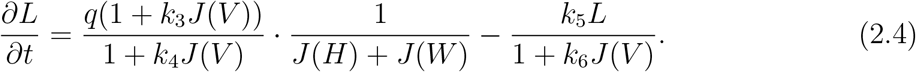

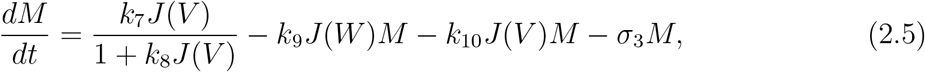

Here *H* is the concentration of uninfected hepatocytes, *W* of infected hepatocytes, *V* virus, *M* ensemble of factors of immune response (space independent), *L* intracellular lipids. This system of equations is considered on the whole real axis in the mathematical analysis presented below or on a bounded and sufficiently long interval with appropriate boundary conditions in numerical simulations. In the case of the whole axis, we use notation

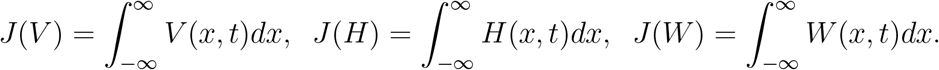

The integrals are taken with respect to the space interval if it is considered to be bounded.

The first term in the right-hand side of equation (2.1) describes proliferation of hepatocytes. The proliferation rate is limited by the maximal concentration *H*_0_ (density-dependent proliferation) meaning that the cell concentration cannot exceed some given value. The second term in this equation characterizes hepatocyte infection by HCV virus.

The next equation contains the same term with sign plus followed by the term corresponding to the elimination of infected cells by factors of immune response, while the last term represents the death rate of infected cells.

Equation (2.3) for the virus concentration contains the diffusion term describing random virus motion, the replication term, virus elimination by the immune factors and virus death rate independent of the immune response. Note that the virus replication rate by infected cells depends on the lipid concentration *L* inside hepatocytes. Increase of intracellular lipid concentration promotes virus replication (*p*_1_ *> p*_2_).

The first term in the right-hand side of equation (2.4) for the intracellular lipid concentration *L* describes its influx from the adipocyte tissue. It is up-regulated by virus through insulin resistance (*k*_3_ *> k*_4_). The total influx is recalculated with respect to one cell due to the term *J* (*H*) + *J* (*W* ) in the denominator. Lipid consumption and transport are down-regulated through inhibition of beta-oxidation of fatty acids and other mechanisms.

We consider in this model a simplified representation of the immune response by the ensemble of immune factors *M* which include innate and adaptive parts with the corresponding cells (macrophages, dendritic cells, T- and B-lymphocytes), molecules (interferon, cytokines), and antibodies. In this simplifies description, *M* eliminate infected cells (equation (2.2)) and viral particles (equation (2.3)). Production of immune factors in equation (2.5) is stimulated by virus antigens proportional to *J* (*V* ). This term is considered with saturation with respect to the amount of antigen taking into account a limited capacity of the immune response.The next two terms describe depletion of the immune factors in the interaction with infected cells and virus particles. The last term characterizes degradation of the immune factors.

## 3 ODE version of the model

We develop an ordinary differential equation (ODE) version of the system described by equation (2.1)-equation (2.5) in order to analyse the system’s dynamical behavior. In particular, transforming the model into its ODE formulation allows us to investigate the existence and stability of equilibrium solutions, as well as to examine how the system responds to perturbations under varying parameter regimes.

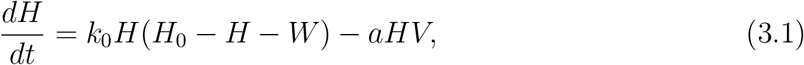

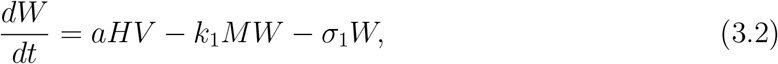

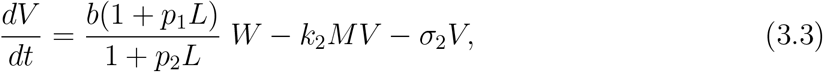

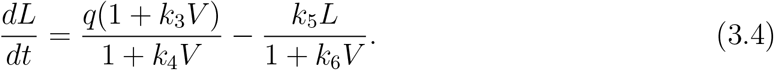

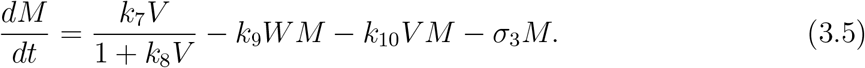

In order to find analytically the stationary points of this system, we simplify the model assuming that immune response is proportional to the viral load *V* without saturation (*k*_8_ = 0) and that cells of the immune response do not deplete in their interaction with infected cells and free virions (*k*_9_ = *k*_10_ = 0). These processes are not so essential for the infection progression and do not change its qualitative dynamics.

### 3.1 Quasi Steady State of the Lipid Variable

Intracellular lipid pools (triglyceride synthesis, *β*-oxidation, lipid-droplet turnover) equilibrate on the scale of hours, whereas hepatocyte infection, immune clearance and net viral load evolve on the scale of days to weeks. Formally, introduce the small parameter

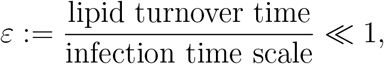

and rescale (3.4) as

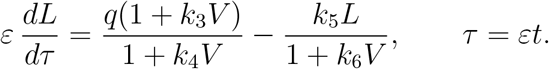

In the singular limit *ε* → 0 (fast lipid relaxation), the lipid variable is slaved to whatever viral load *V* happens to be at that instant: we obtain

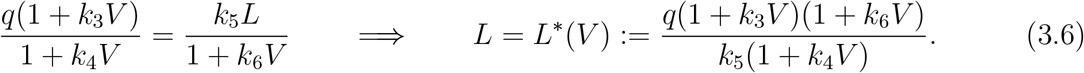

We applied the standard singular-perturbation (Tikhonov) argument for eliminating a fast variable from a slow–fast system: provided the “boundary layer” system *dL/dτ* = *q*(1 + *k*_3_*V* )*/*(1 + *k*_4_*V*) − *k*_5_*L/*(1 + *k*_6_*V* ) (with *V* frozen) has *L*^*∗*^(*V* ) as a globally attracting equilibrium which holds here since the right-hand side is affine-decreasing in *L* with negative slope −*k*_5_*/*(1+*k*_6_*V* ) *<* 0 for all *V* ≥ 0 – the reduced (slow) dynamics obtained by substituting *L* = *L*^*∗*^(*V* ) into (3.3) approximate the full system uniformly on compact time intervals, and *exactly* reproduce every stationary point of the full five-dimensional system, since at any true equilibrium *dL/dt* = 0 automatically forces *L* = *L*^*∗*^(*V* ) regardless of *ε*.

#### Lemma 1

(Monotonicity of the lipid profile). *If k*_3_ + *k*_6_ ≥ *k*_4_, *then L*^*∗*^(*V* ) *is nondecreasing in V for V* ≥ 0, *with* 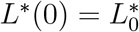. *If in addition p*_1_ ≥ *p*_2_, *then* Φ(*L*^*∗*^(*V* )) *is nondecreasing in V for V* ≥ 0 *where*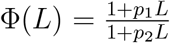.

*Proof*. Write 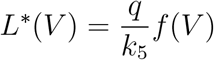with 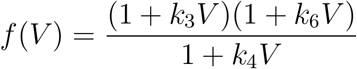. Then

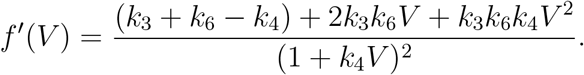

Since *k*_3_, *k*_4_, *k*_6_ ≥ 0, both 2*k*_3_*k*_6_*V* and *k*_3_*k*_6_*k*_4_*V* ^2^ are nonnegative for *V* ≥ 0, so *f* ^*′*^(*V* ) ≥ 0 whenever *k*_3_ + *k*_6_ ≥ *k*_4_. This proves the first claim. For the second, if 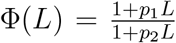 then Φ^*′*^(*L*) = (*p*_1_ − *p*_2_)*/*(1 + *p*_2_*L*)^2^ ≥ 0 when *p*_1_ ≥ *p*_2_, so Φ is nondecreasing; composing with the nondecreasing map *V* ↦ *L*^*∗*^(*V* ) gives the result. □

Substituting (3.6) into Φ(*L*) defines the *lipid-corrected effective production rate*

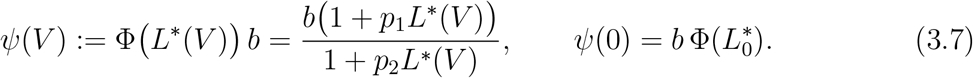

Equations (3.1), (3.2), (3.5), together with (3.3) rewritten as

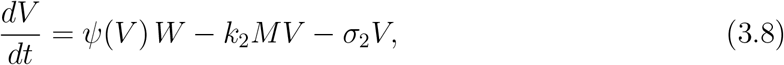

now form a closed four-dimensional system (*H, W, V, M* ) in which all lipid dependence enters only through the scalar function *ψ*(*V* ).

### 3.2 Stationary points

We seek stationary points with *H* ≠ 0 (the biologically relevant case). We obtain the following (with *k*_8_ = 0),

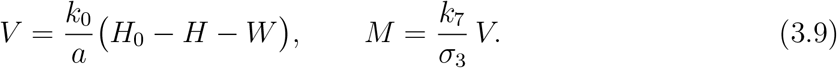

Introduce, exactly as in the lipid-free analysis,

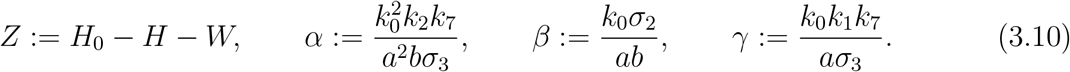

so that *V* = (*k*_0_*/a*)*Z* and *M* = (*k*_0_*k*_7_*/aσ*_3_)*Z*.

Substituting (3.9) into the stationary form of the reduced virus equation (3.8),

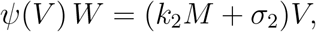

and using (3.10), gives

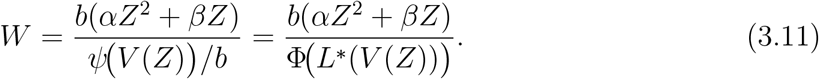

When Φ ≡ 1 (i.e. *p*_1_ = *p*_2_ = 0), (3.11) reduces exactly to the lipid-free relation *W* = *αZ*^2^+*βZ* used in the base model.

Substituting (3.9) into the stationary form of (4.8),

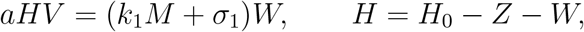

and using (3.9) again yields

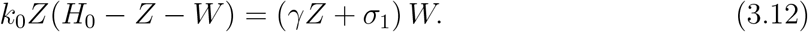

Combining (3.11) and (3.12) and dividing through by *Z* (the branch *Z* = 0 is treated separately below) gives a single scalar equation for the stationary excursion *Z* = *H*_0_ −*H* −*W* :

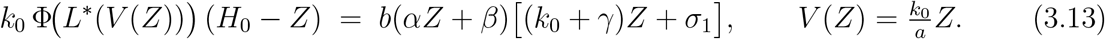

Define

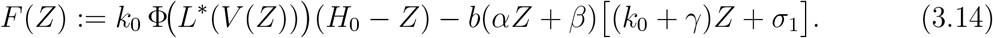

Every positive root of *F* (*Z*) = 0 on (0, *H*_0_) corresponds, through (3.9) and (3.11), to an endemic stationary point (*H*^*∗*^, *W* ^*∗*^, *V* ^*∗*^, *M* ^*∗*^).

*Z* = 0 always solves (3.13). Then *V* = 0, hence *M* = 0 (from (3.9)), and (3.12) forces *W* = 0. Consequently *H* = *H*_0_. Hence the disease-free equilibrium

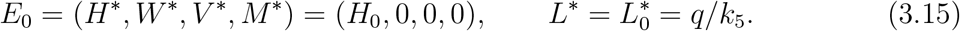

Note that this equilibrium does not depend on *p*_1_, *p*_2_: in the absence of virus and infected cells, the lipid feedback term never enters the dynamics.

#### Proposition 1

(Threshold for endemic equilibrium). *Define the lipid-induced basic reproduction number*

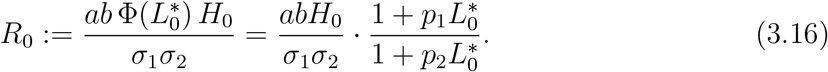

*If R*_0_ *>* 1, *equation* (3.13) *has at least one solution Z*^*∗*^ ∈ (0, *H*_0_), *and hence at least one endemic equilibrium exists. If R*_0_ *<* 1 *and* Φ *is nondecreasing (Lemma 1), no positive endemic equilibrium exists near Z* = 0, *and E*_0_ *is the only biologically admissible equilibrium in a neighbourhood of the origin*.

*Proof. F* defined in (3.14) is continuous on [0, *H*_0_]. At *Z* = 0, *V* (0) = 0, 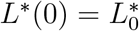, so

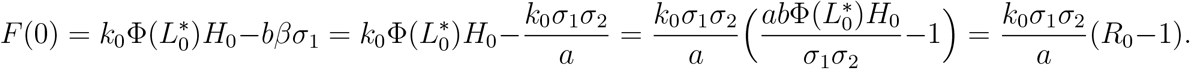

Hence *F* (0) *>* 0 whenever *R*_0_ *>* 1. At *Z* = *H*_0_ (i.e. *H*^*∗*^ = 0, the boundary of the biologically admissible region), the first term of (3.14) vanishes while the second term is strictly negative for any admissible parameter values, so *F* (*H*_0_) *<* 0. By the intermediate value theorem, *F* has at least one zero in (0, *H*_0_) when *R*_0_ *>* 1. Conversely, if *R*_0_ *<* 1 then *F* (0) *<* 0; since Φ(*L*^*∗*^(*V* (*Z*))) is nondecreasing (Lemma 1) while (*H*_0_ − *Z*) is decreasing, and the right-hand side *b*(*αZ* + *β*)[(*k*_0_ + *γ*)*Z* + *σ*_1_] is increasing and convex in *Z, F* remains negative for small *Z >* 0, so no small positive root exists.

In the lipid-free case Φ≡ 1, the left-hand side *k*_0_(*H*_0_− *Z*) of (3.14) is strictly decreasing in *Z* while the right-hand side is strictly increasing, so *at most one* positive root exists. With lipid feedback, Φ(*L*^*∗*^(*V* (*Z*))) is nondecreasing in *Z* under the hypotheses of Lemma 1, so the left-hand side of (3.13) is the product of a decreasing factor (*H*_0_−*Z*) and a nondecreasing factor Φ(*L*^*∗*^(*V* (*Z*))); its monotonicity is no longer guaranteed in general. Consequently (3.13) may in principle admit more than one positive root, i.e. the lipid feedback can in general support multiple endemic equilibria for a fixed parameter set, even though every numerical continuation reported for this model exhibits a single branch. A sufficient condition ruling out multiplicity is that Φ(*L*^*∗*^(*V* (*Z*)))(*H*_0_ − *Z*) remains strictly decreasing on (0, *H*_0_), which holds automatically when *p*_1_ is small or *L*^*∗*^ saturates quickly in *V* .

The explicit form of the endemic equilibrium for any root *Z*^*∗*^ of (3.13),

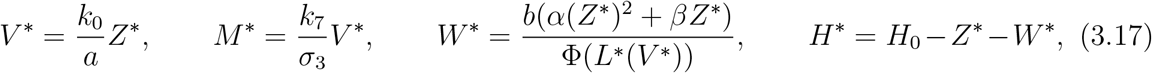

and the corresponding stationary lipid level follows from (3.6),

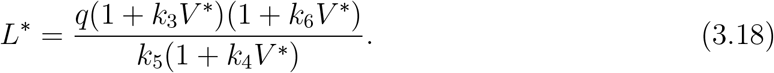

### 3.3 Local stability analysis

We linearize the reduced four-dimensional system (3.1), (3.2), (3.8), (3.5) about a stationary point (*H*^*∗*^, *W* ^*∗*^, *V* ^*∗*^, *M* ^*∗*^), treating *L* as slaved to *V* through equation (3.6).

The only structural difference from the lipid-free Jacobian is that the production term in (3.8) is *ψ*(*V* )*W* = *b* Φ(*L*^*∗*^(*V* )) *W* rather than *bW*, so its partial derivative with respect to *V* acquires an extra chain-rule contribution,

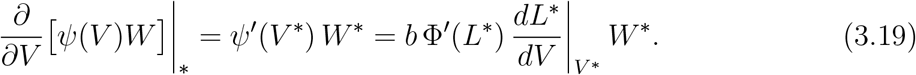

Write *κ* := *ψ*^*′*^(*V* ^*∗*^)*W* ^*∗*^ ≥0 ; by Lemma 2, *κ* ≥0 whenever *p*_1_ ≥*p*_2_ and *k*_3_ + *k*_6_ ≥*k*_4_.

The Jacobian is

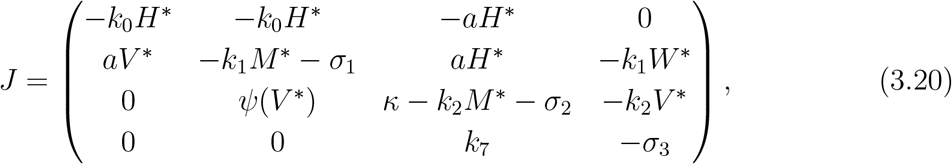

where the (1, 1)-entry uses the stationary relation *k*_0_(*H*_0_ −*H*^*∗*^ −*W* ^*∗*^) = *aV* ^*∗*^ to eliminate the (vanishing) first-order term.

#### 3.3.1 Stability of the disease-free equilibrium

At *E*_0_ = (*H*_0_, 0, 0, 0), *W* ^*∗*^ = 0, so *κ* = *ψ*^*′*^(0) · 0 = 0 vanishes identically: *the lipid feedback does not affect the linear stability of the disease-free equilibrium*. The characteristic equation reduces, exactly as in the lipid-free analysis, to

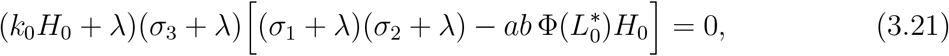

with 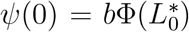. The bracketed factor has a positive root if and only if 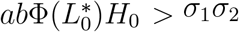, i.e. if and only if *R*_0_ *>* 1 (cf. (3.16)).

##### Corollary 1.

*E*_0_ *is locally asymptotically stable if R*_0_ *<* 1 *and unstable if R*_0_ *>* 1, *where R*_0_ *is the reproduction number* (3.16). *A nonzero baseline hepatic lipid level* 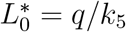 *together with p*_1_ *> p*_2_ *strictly lowers the clearance threshold relative to the lipid-free model, since* 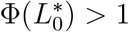 *in that case*.

#### 3.3.2 Stability of the endemic equilibrium

At an endemic equilibrium (*W* ^*∗*^ *>* 0), the characteristic polynomial det(*J* − *λI*) = 0 is a quartic

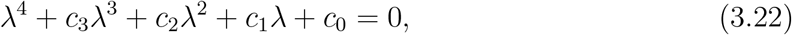

whose coefficients, obtained by expanding (3.20), are

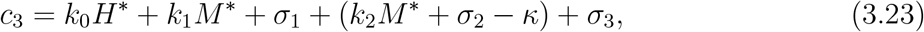

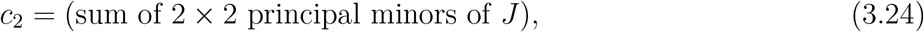

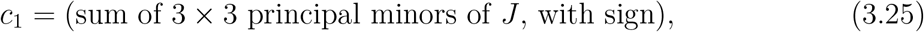

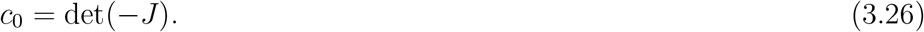

The explicit expressions for *c*_2_, *c*_1_, *c*_0_ are lengthy but are all *linear* in the single new parameter *κ*, since *κ* enters *J* in only one entry. Writing *c*_*i*_(*κ*) = *c*_*i*_(0) + *κ d*_*i*_ for *i* = 0, 1, 2, 3, where *c*_*i*_(0) are exactly the lipid-free coefficients obtained by setting *p*_1_ = *p*_2_ = 0 (equivalently *κ* = 0), one finds

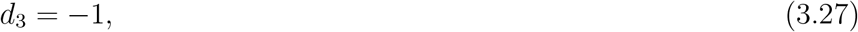

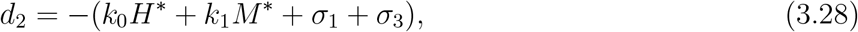

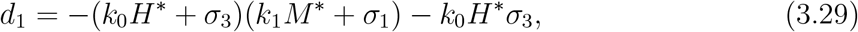

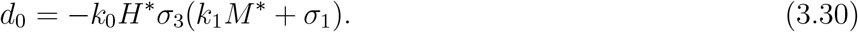

All *d*_*i*_ ≤ 0, so *increasing κ decreases every coefficient c*_*i*_ *of the characteristic polynomial*. By the Routh–Hurwitz criterion, local asymptotic stability of a quartic requires

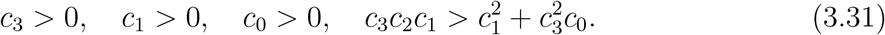

Since *κ* enters every coefficient with a nonpositive sign, increasing *κ* (equivalently, increasing the lipid amplification parameter *p*_1_, since 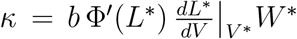 is increasing in *p*_1_ by Lemma 2) monotonically erodes the Routh–Hurwitz margins in (3.31): it decreases (reducing the net damping of the system, i.e. −trace(*J* )), and decreases *c*_1_, *c*_0_ while entering the Hurwitz determinant 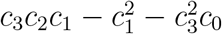 non-monotonically through cross terms. In the regime where *c*_3_ is the first coefficient to lose positivity, the mechanism is a classical *Hopf-type* destabilization: a complex-conjugate pair of eigenvalues crosses the imaginary axis as *κ* increases through a critical value *κ*_*H*_, producing sustained oscillations, while for *κ* even larger the equilibrium can lose stability outright through a real eigenvalue crossing (loss of positivity of *c*_0_).

##### Proposition 2

(Destabilizing role of lipid feedback). *At a fixed endemic equilibrium with W* ^*∗*^ *>* 0, *the local stability margin of the endemic equilibrium is a nonincreasing function of κ* = *ψ*^*′*^(*V* ^*∗*^)*W* ^*∗*^. *In particular, for p*_1_ ≥ *p*_2_ *and k*_3_ + *k*_6_ ≥ *k*_4_ *(so that κ* ≥ 0 *and κ is nondecreasing in p*_1_*), sufficiently strong lipid amplification destabilizes an endemic equilibrium that would be stable in the lipid-free limit (p*_1_ = 0*), producing a Hopf bifurcation into sustained oscillations*.

This is exactly the destabilization mechanism observed numerically in the model: for fixed *R*_0_ obtained by tuning *a*, increasing *p*_1_ moves the system from a stable monotone or damped-oscillatory endemic state into a sustained oscillatory (limit-cycle) regime once *p*_1_ exceeds a critical value (numerically *p*_1_ ≈ 0.337 in the reference parameterization), while the disease-free threshold itself, governed only by 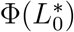 through (3.21), is comparatively insensitive to *p*_1_ over the same range, since 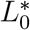 is independent of *p*_1_ and only the (typically small) baseline value 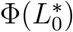 enters the disease-free characteristic equation. This shows precisely how lipid accumulation enters both the threshold 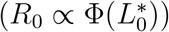 and, more importantly, the *stability* of the endemic state through the extra Jacobian term *κ* in (3.20). Because *κ* is proportional to the local slope of the lipid amplification factor evaluated at the endemic viral load, and this slope grows with *p*_1_, the analysis identifies lipid accumulation as an intrinsically destabilizing positive feedback on the endemic state, which analytically explains, the transition from stable to oscillatory and ultimately relapsing viral dynamics observed as *p*_1_ (or, equivalently, *b*) is increased in numerical simulations of the model.

#### 3.3.3 Lipid feedback lowers the bifurcation threshold: a transcritical bifurcation at *R*_0_ = 1

The endemic branch bifurcates from the disease-free equilibrium exactly at *R*_0_ = 1, where

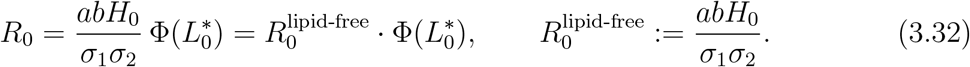

Because 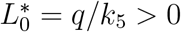 is the (nonzero) pre-infection hepatic lipid level, and 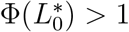 whenever *p*_1_ *> p*_2_, the lipid-corrected threshold is crossed *earlier* than the purely viral threshold: a baseline hepatic lipid environment alone (independent of any infection-induced lipid accumulation) is enough to lower the effective clearance bar. This is the precise mechanistic content of the “lipid accumulation is not merely a consequence but a driver” claim: even the pre-existing lipid economy of the hepatocyte enters the invasion threshold multiplicatively.

Near the threshold, write *r* := *R*_0_ − 1 as a small bifurcation parameter. From the proof of Proposition 1,

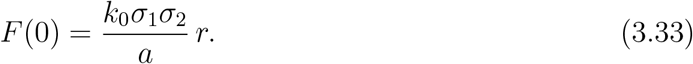

Differentiating the master equation (3.14) at *Z* = 0 (using *V* (0) = 0, 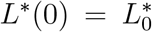, and *V* ^*′*^(*Z*) = *k*_0_*/a*) gives

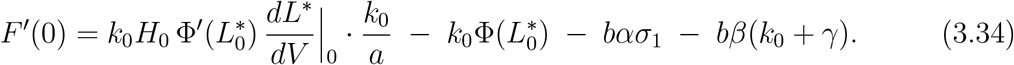

The first term in (3.34) is the direct signature of the lipid feedback on the bifurcation itself: it is nonnegative whenever *p*_1_ ≥ *p*_2_ and *k*_3_ + *k*_6_ ≥ *k*_4_ (Lemma 2), and it partially offsets the otherwise strictly negative remaining terms. Provided *F* ^*′*^(0) = 0, the generic case, the implicit function theorem gives a locally unique branch

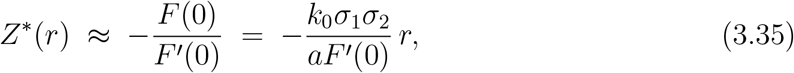

which is positive for *r >* 0 whenever *F* ^*′*^(0) *<* 0: this is the classical *transcritical exchange of stability* between *E*_0_ and *E*^*∗*^ familiar from single-strain infection models, here generalized so that its slope (3.35), and hence how rapidly the endemic viral load and infected-cell population grow just above threshold, is itself modulated by the lipid-feedback correction term in (3.34). A larger lipid sensitivity 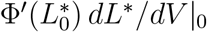flattens *F* ^*′*^(0) toward zero, which *steepens* the emergent branch *Z*^*∗*^(*r*): lipid feedback not only lowers the threshold but makes the post-threshold rise of the endemic state sharper.

#### 3.3.4 The endemic state is intrinsically hyperlipidemic

##### Corollary 2

(Elevated stationary lipid level at the endemic state). *Suppose k*_3_ + *k*_6_ ≥ *k*_4_. *Then at any endemic equilibrium* 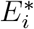 *with* 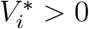,

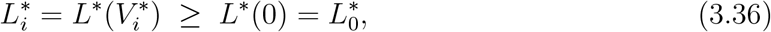

*with strict inequality whenever k*_3_ + *k*_6_ *> k*_4_. *That is*, every *endemic equilibrium of the model carries a hepatocyte lipid level at or above the pre-infection baseline; the endemic state is intrinsically hyperlipidemic relative to the disease-free state*.

*Proof*. Immediate from the monotonicity of *L*^*∗*^(*V* ) in *V* and 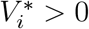. □

Corollary 2 is the equilibrium-level counterpart of the biological hypothesis of the model, that chronic HCV infection at steady state cannot coexist with a disease-free hepatic lipid profile. Steatosis is not an optional comorbidity that may or may not accompany the endemic state, it is a structural property of every endemic equilibrium admitted by the model, exactly mirroring the clinical observation that hepatic steatosis is detected in a large fraction of chronic hepatitis C patients and correlates with viral load.

#### 3.3.5 Self-consistency: the endemic equilibrium is a fixed point of a feedback loop

The master equation (3.14) makes the feedback structure explicit: Φ(*L*^*∗*^(*V* (*Z*))) appears evaluated at the *same Z* that the equation is solving for. Biologically, this encodes a closed loop:

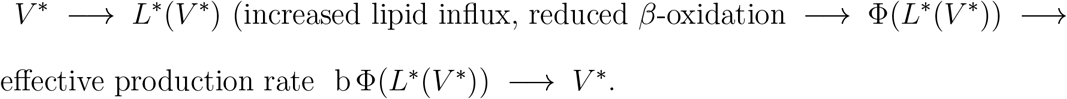

The endemic equilibrium 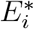 is precisely the fixed point of this loop: a viral load that, through the induced lipid accumulation, sustains exactly the level of viral production needed to reproduce that same viral load against clearance. This is a positive feedback loop in the classical sense, and it is the mechanism responsible for i) amplification: comparing two models with the same viral and immune kinetic parameters (*a, b, σ*_1_, *σ*_2_, *k*_1_, *k*_2_, *k*_7_) but Φ ≡ 1 (lipid-free) versus Φ ≢ 1 with *p*_1_ *> p*_2_, the lipid-aware model has a strictly larger effective production rate at any positive viral load, *b*Φ(*L*^*∗*^(*V* )) *> b* for *V >* 0; since *F* is increasing in the coefficient multiplying Φ inside (3.14), the lipid-aware endemic root *Z*^*∗*^ is pushed to a larger value than its lipid-free counterpart for the same kinetic parameters consistent with the numerically observed rightward/upward shift of the endemic viral load as *p*_1_ increases; ii) persistence: by Proposition 1, the enlarged effective production rate can sustain *R*_0_ *>* 1 in parameter regions where a lipid-free model with the same *a, b, H*_0_, *σ*_1_, *σ*_2_ would predict clearance 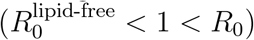, i.e. lipid feedback can convert a self-limiting infection into a chronic one purely through the multiplicative factor 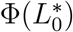, without any change in the viral kinetic parameters themselves.

This produces a single coherent narrative on existence and stability:

| Regime | Character of $E_i^*$ |
| --- | --- |
| $R_0$ slightly above 1 | $Z_i^*$ small (by (3.35)), so $V_i^*$ small, so $L_i^* \approx L_0^*$ and $\kappa \approx 0$: the endemic equilibrium is stable, approached monotonically. |
| $R_0$ moderately above 1 | $V_i^*$, hence $L_i^*$ and $\kappa$, grow; Routh–Hurwitz margins shrink: the approach to $E_i^*$ becomes a damped spiral (complex eigenvalues with negative real part). |
| $R_0$ (or $p_1$) large | $\kappa$ exceeds the critical value at which a Routh–Hurwitz condition in (3.31) fails: $E_i^*$ loses stability through a Hopf bifurcation, and the system settles onto a sustained limit cycle (relapsing viral dynamics). |

This table is the analytical counterpart of the four qualitative regimes (clearance, stable monotone endemic, damped-oscillatory endemic, sustained oscillatory): all four are organized by the single scalar quantity *κ*, which is itself entirely determined by how strongly, and how sensitively, the endemic lipid level departs from baseline.

### 3.4 Dynamical regimes of the ODE model

We first consider representative ODE parameter sets chosen to display the main qualitative regimes. The simulations confirm the threshold behavior predicted by the analytical study. For *R*_0_ *<* 1, the disease-free equilibrium is stable. In the representative case *R*_0_ = 0.97, the viral load initially increases and reaches a large transient peak, but subsequently decreases to values below the clearance threshold. Thus, even when a substantial transient infection occurs, the long-time behavior is elimination of infection.

For *R*_0_ *>* 1, a positive endemic equilibrium appears. Close to the threshold, this equilibrium is stable and approached monotonically after an initial overshoot. For larger values of *R*_0_, the approach to equilibrium becomes oscillatory. Finally, for sufficiently large *R*_0_, the endemic equilibrium loses stability and the solution approaches a sustained oscillatory regime (Figure 2).

**Figure 2:**
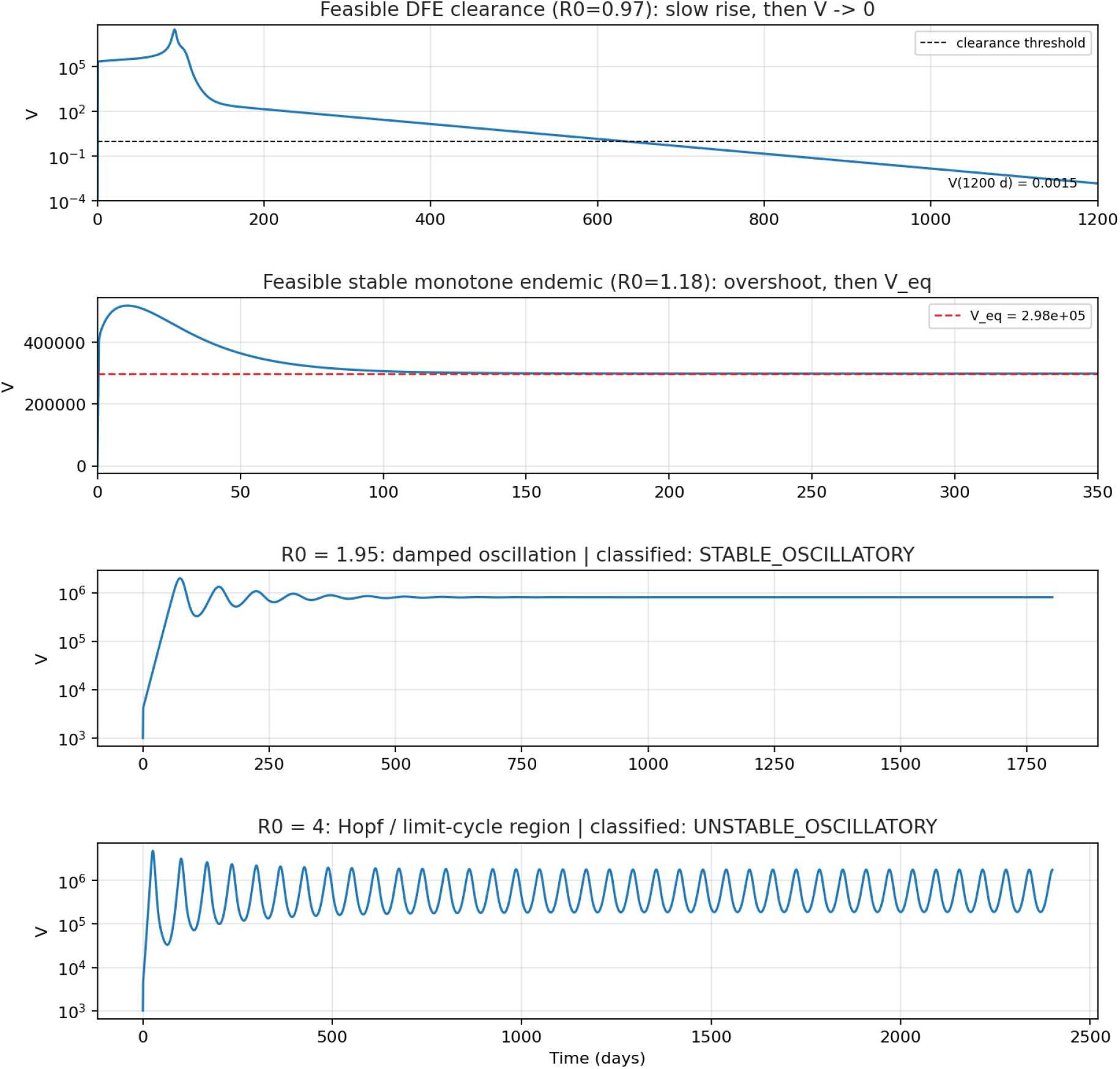
Representative viral-load trajectories for four qualitative regimes of the ODE model. For *R*_0_ = 0.97, the solution exhibits a large transient peak followed by clearance. For *R*_0_ = 1.18, the viral load converges monotonically to a positive endemic equilibrium after an initial overshoot. For *R*_0_ = 1.95, the endemic equilibrium is approached through damped oscillations. For *R*_0_ = 4, the solution remains in a persistent oscillatory regime.

The corresponding threshold diagram is shown in Figure 3. The disease-free equilibrium loses stability at *R*_0_ = 1. A stable monotone endemic regime is observed immediately above this threshold, followed near *R*_0_ ≈ 1.21 by a stable endemic equilibrium that is approached through damped oscillations. A further transition occurs near *R*_0_ ≈ 2.78, where the leading eigenvalue crosses the imaginary axis and the endemic equilibrium becomes unstable. These simulations indicate that the model can reproduce acute clearance, stable chronic infection, damped recurrent infection, and sustained relapsing dynamics within the same model structure.

**Figure 3:**
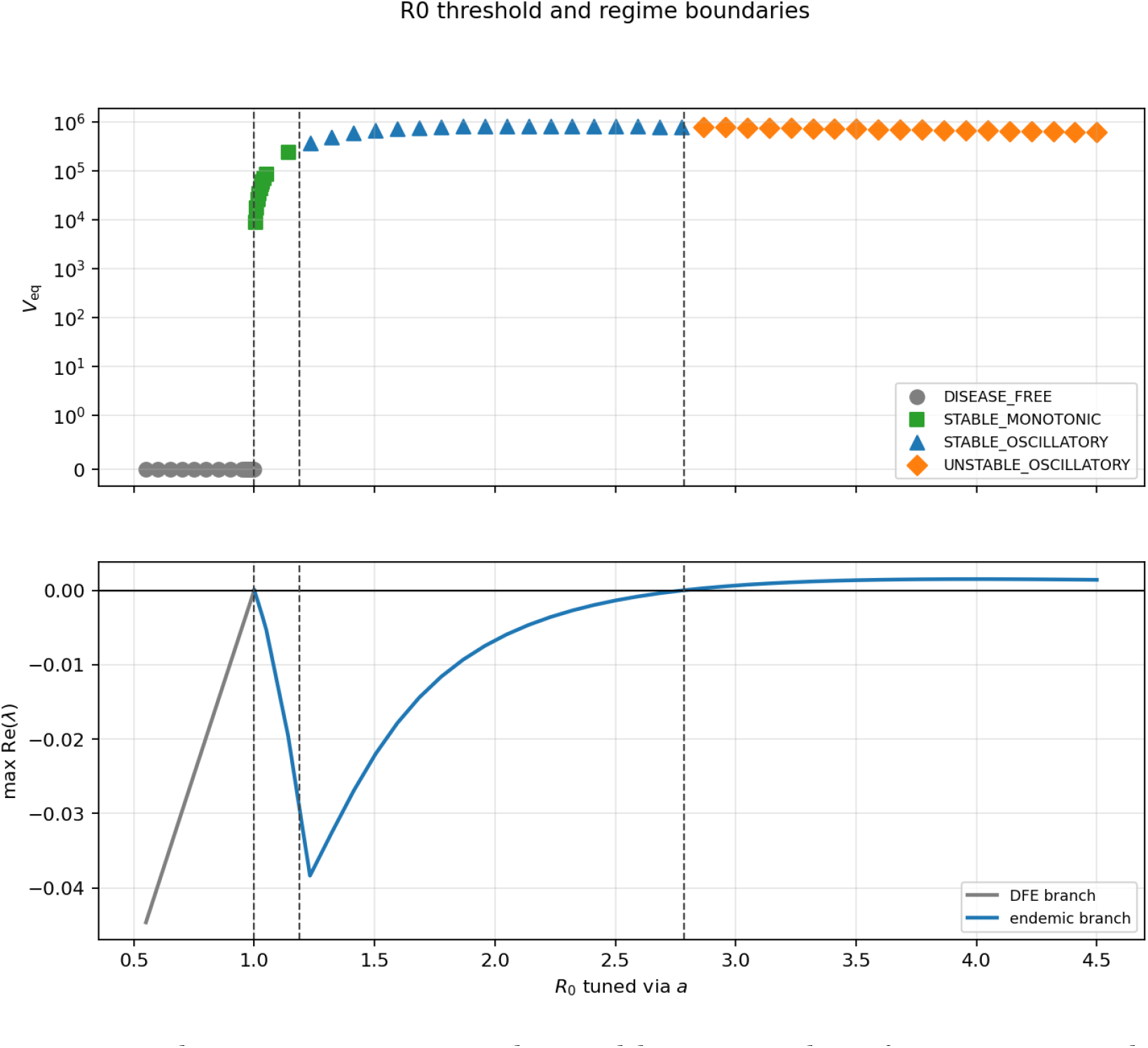
Numerical continuation in *R*_0_, obtained by tuning the infection rate *a*. The upper panel shows the equilibrium viral load, classified by linear stability. The lower panel shows the largest real part of the active eigenvalues. The disease-free threshold occurs at *R*_0_ = 1, followed by transitions to oscillatory convergence and then to an unstable oscillatory endemic equilibrium.

The lower panel of Figure 3 should be interpreted as a numerical stability diagnostic rather than as a single smooth eigenvalue branch. At each sampled value of *R*_0_, the equilibrium is reconstructed from a finite-time trajectory and the plotted quantity is the largest real part among the active eigenvalues. This maximum can switch from one eigenvalue branch to another, especially near the monotone oscillatory transition where a complex pair becomes relevant. In addition, the plotted branch changes from the disease-free equilibrium below threshold to an endemic equilibrium above threshold. These branch and equilibrium switches explain the lack of visual smoothness in the reported eigenvalue curve.

#### 3.4.1 Bifurcation

The one-parameter continuation in Figure 3 is in fact a single horizontal slice, at one fixed lipid sensitivity *p*_1_, through a two-parameter bifurcation structure. Figure 4 summarizes this structure in the (*R*_0_, *p*_1_) plane, obtained by two-parameter numerical continuation of the reduced ODE system.

**Figure 4:**
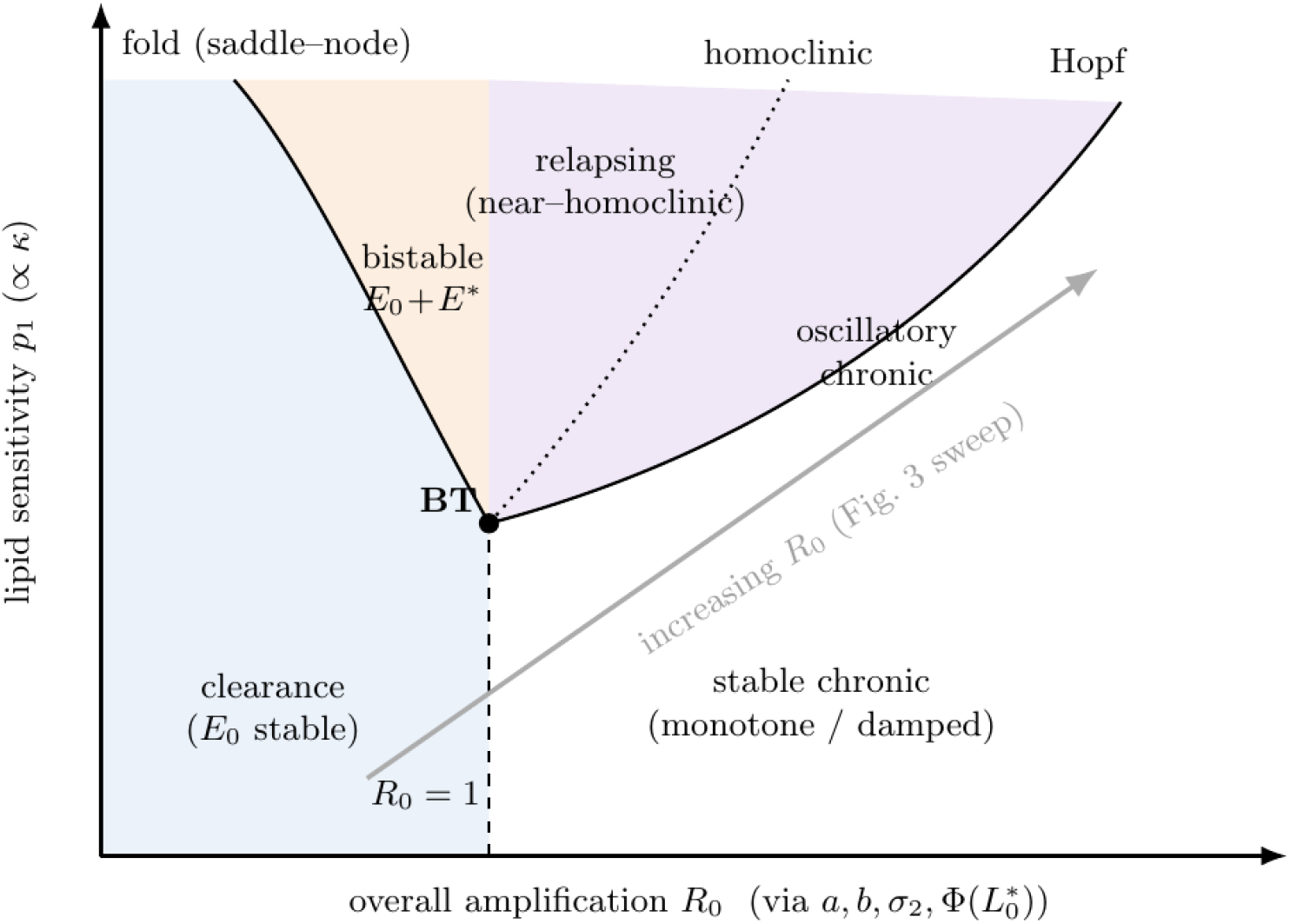
Two-parameter bifurcation diagram in the (*R*_0_, *p*_1_) plane. The fold (saddle-node) curve and the transcritical line *R*_0_ = 1 meet at a Bogdanov–Takens point (BT), separating the clearance region (*E*_0_ stable) from a narrow bistable wedge (*E*_0_ + *E*^*∗*^ both stable) and from the stable chronic region (monotone or damped convergence to *E*^*∗*^). A Hopf curve emanating from BT bounds the oscillatory chronic region; a nearby homoclinic curve bounds a thin relapsing (near-homoclinic) wedge between the Hopf and homoclinic curves. The gray arrow indicates the direction of the *R*_0_-only sweep reported in Figure 3.

For small *p*_1_, the disease-free equilibrium *E*_0_ loses stability exactly at *R*_0_ = 1 through the transcritical bifurcation established in (Corollary 1), and the emergent endemic branch *E*^*∗*^ is initially stable, consistent with Section 3.3.2. As *p*_1_ increases, however, the destabilizing lipid-feedback term *κ* in eqs. (3.19)-(3.20) grows, and the transcritical bifurcation at *R*_0_ = 1 becomes subcritical: a saddle-node (fold) curve peels away from the line *R*_0_ = 1 at a Bogdanov–Takens (BT) point which is a codimension-2 bifurcation point of a two-dimensional (or higher) dynamical system, where the equilibrium has a double zero eigen-value and in our case it bends into the region *R*_0_ *<* 1, bounding a narrow bistable wedge in which the disease-free state and a stable endemic equilibrium coexist (*E*_0_ + *E*^*∗*^). Biologically, in this wedge a sufficiently large initial viral inoculum can establish chronic infection even though the linearized invasion threshold *R*_0_ *<* 1 would, by (Corollary 1), otherwise predict clearance, a hysteresis effect due entirely to the hepatic lipid feedback loop identified in Section 3.3.5, and absent in the lipid-free limit of small *p*_1_.

For *p*_1_ above the BT value, the endemic branch is stable near onset but loses stability through the Hopf mechanism of Proposition 2 as *R*_0_ increases further, producing the Hopf curve that bounds the “oscillatory chronic” region. A homoclinic curve emanates from the same BT point, running just above the Hopf curve; between the two lies a thin “relapsing” wedge in which the limit cycle born at the Hopf bifurcation grows rapidly in period and amplitude as it approaches a homoclinic connection, producing the long, near-silent inter-episode intervals characteristic of clinically relapsing HCV viremia, as opposed to the more regular, moderate-amplitude oscillations found deeper inside the oscillatory-chronic region. The gray arrow schematically indicates the direction of the *R*_0_-sweep already reported in Figure 3, its two coarse regime transitions, clearance to chronic infection near *R*_0_ = 1 and stable to sustained-oscillatory chronic infection near *R*_0_ ≈ 2.78, correspond respectively to crossing the transcritical/fold boundary and the Hopf curve in Figure 4. The finer monotone-to-damped-oscillatory transition reported near *R*_0_ ≈ 1.21 occurs within the single “stable chronic” region here and is not separately delineated by a bifurcation curve at this coarser, two-parameter resolution.

### 3.5 Dependence on viral and lipid parameters

We varied parameters that directly affect the effective replication capacity of the virus. Increasing the viral production rate *b* increases *R*_0_ and moves the system from clearance to persistent infection and then to large-amplitude oscillations. For small *b*, the viral load decays to zero; for intermediate *b*, infection persists at an endemic level; and for large *b*, recurrent peaks of viral load are observed together with strong depletion and partial recovery of the uninfected hepatocyte population.

A similar transition occurs when the death rate of infected cells, *σ*_1_, is varied. Since *R*_0_ is inversely proportional to *σ*_1_, decreasing *σ*_1_ promotes chronic infection and oscillatory behavior, whereas increasing *σ*_1_ favors clearance. Thus, efficient loss or removal of infected cells stabilizes the disease-free state (Figure 5).

**Figure 5:**
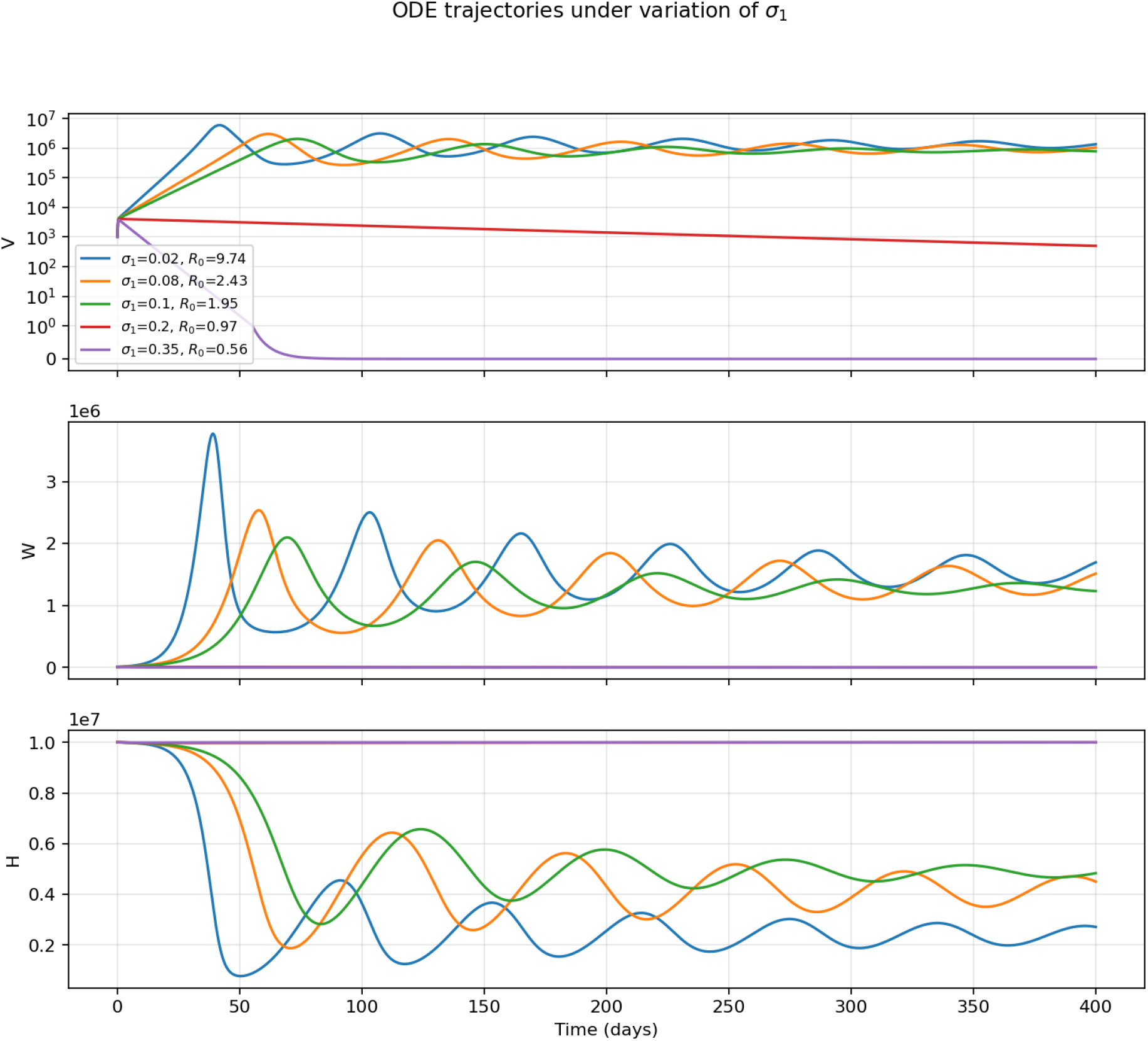
Effect of varying the infected-cell death rate *σ*_1_. Smaller values of *σ*_1_ increase *R*_0_ and produce persistent oscillatory infection, while larger values suppress viral growth and lead to clearance.

The lipid feedback parameter *p*_1_ also changes the qualitative dynamics. Larger values of *p*_1_ increase the lipid-dependent viral production term

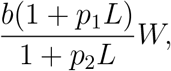

and therefore increase the effective viral amplification. At lower values of *p*_1_, viral load converges to a stable endemic level, while larger values produce oscillatory infection dynamics (Figure 6). This confirms that lipid accumulation can act as a positive feedback on viral replication and can shift the system toward more persistent or recurrent infection. This result demonstrate that lipid accumulation is not merely a consequence of HCV infection but acts as an active driver of viral persistence and recurrent infection through a positive metabolic feedback loop.

**Figure 6:**
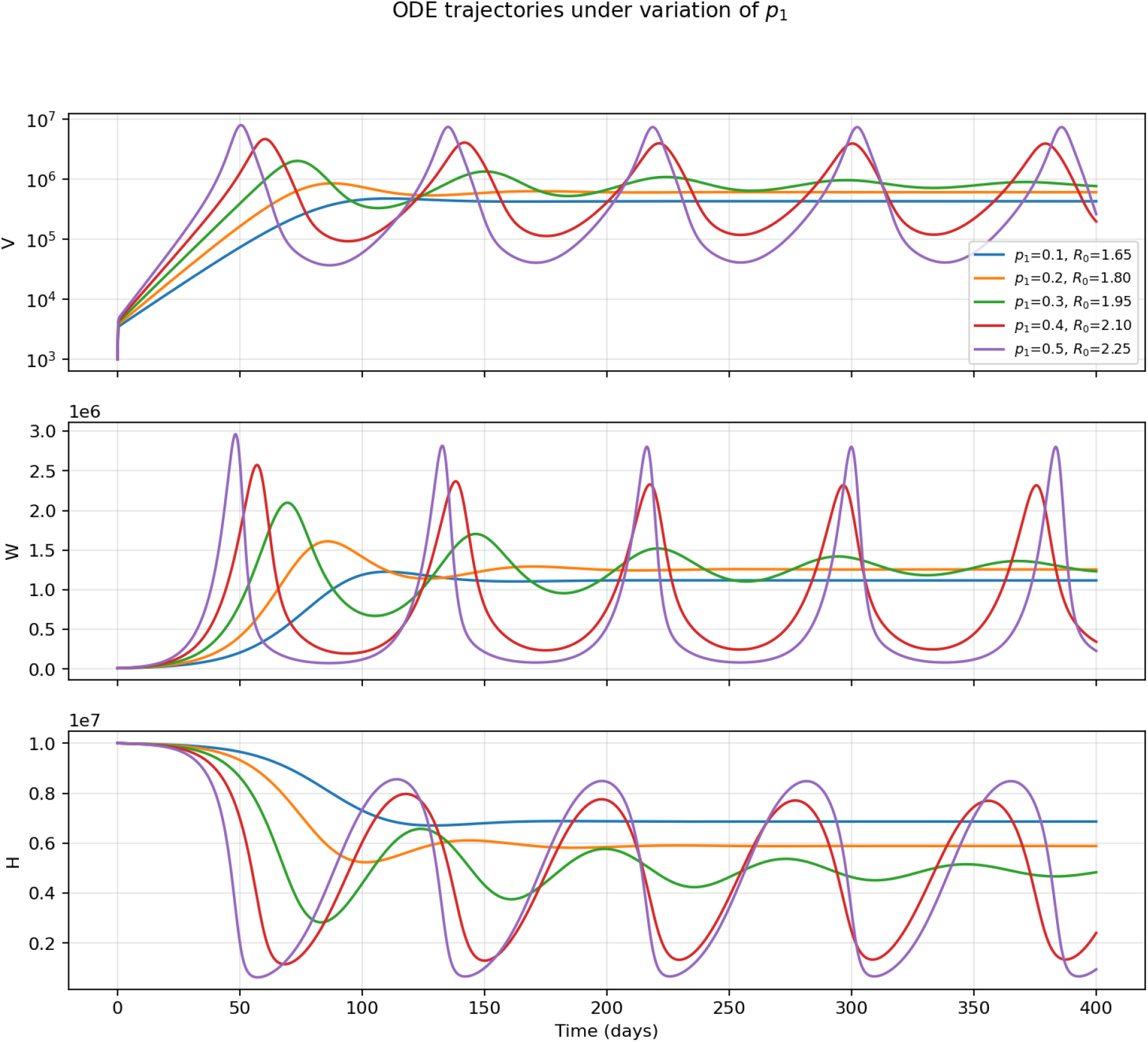
Effect of varying the lipid amplification parameter *p*_1_. Increasing *p*_1_ strengthens lipid-dependent viral production and drives the model from stable endemic infection toward oscillatory recurrent dynamics.

Although increasing b or *p*_1_ increases the effective replication number and produces higher viral peaks, the long term average viral load does not necessarily increase. Once the endemic equilibrium loses stability, the system enters a limit cycle regime characterized by episodic viral outbreaks followed by strong depletion of susceptible hepatocytes and enhanced immune mediated control. Consequently, the virus spends longer periods at low abundance between outbreaks. Biologically, after each outbreak, many hepatocytes become infected, immune response increases,infected cells are removed, healthy hepatocytes fall to very low levels. Since virus production depends on infected cells, and infected cells ultimately depend on available healthy hepatocytes, viral replication collapses after each burst. Therefore, larger *R*_*O*_ values may produce higher peak viral loads while maintaining similar or even lower time averaged viral loads.

To compare these trajectory sweeps with the *R*_0_ threshold diagram, we also computed continuation-style stability diagnostics along the individual parameter directions (Figure 7). These maps show that the numerical boundaries depend on the parameter used to move through the model. Increasing *b* crosses the disease-free threshold near *b* ≈ 1.13, changes from monotone to damped endemic convergence near *b* ≈ 1.31, and loses endemic stability near *b* ≈ 2.35. Increasing *p*_1_ does not cross the disease-free threshold over the plotted range, but it destabilizes the endemic equilibrium near *p*_1_ ≈ 0.337. In contrast, increasing *σ*_1_ moves the system toward clearance, with a damped-to-monotone change near *σ*_1_ ≈ 0.174 and a disease-free transition near *σ*_1_ ≈ 0.196. Increasing viral clearance *σ*_2_ reverses the oscillatory instability near *σ*_2_ ≈ 6.0, approaches monotone endemic convergence near *σ*_2_ ≈10.9, and reaches clearance near *σ*_2_ ≈12.9. The *σ*_2_ row is included as a numerical continuation diagnostic rather than as a purely literature-bounded sweep: the upper part of this scan extends beyond the anchor range 5.5–8.3 day^*−*1^. Thus the transitions near *σ*_2_ ≈10.9 and *σ*_2_ ≈12.9 should be interpreted as model-extrapolation thresholds for this parameter direction.

**Figure 7:**
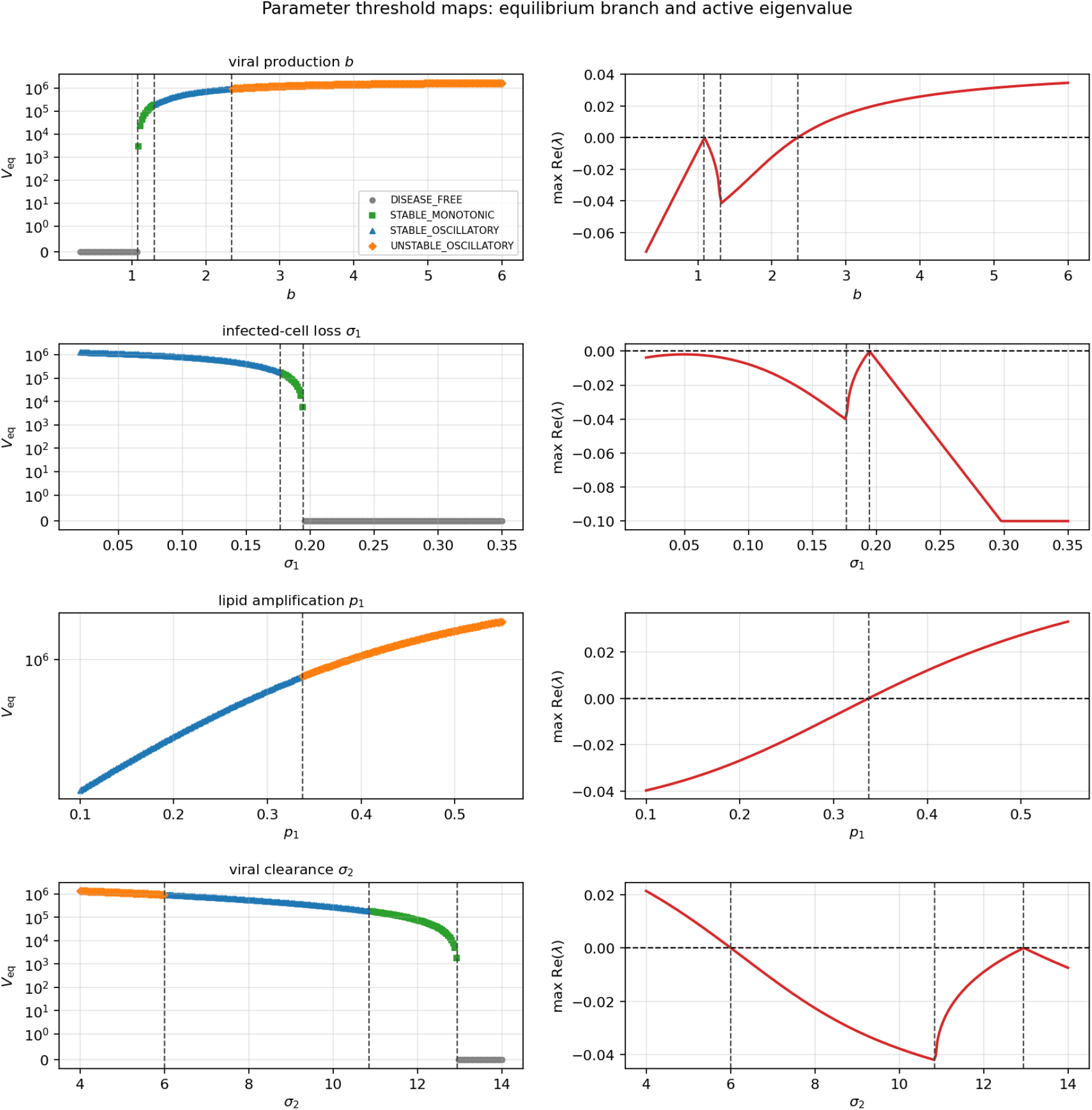
Continuation-style diagnostics for the parameters varied in Figures 5–6, together with viral clearance *σ*_2_. Each row shows the equilibrium viral load classified by linear regime and the leading active real eigenvalue. Vertical dashed lines mark sampled regime changes. These parameter-specific maps show that thresholds found by tuning *R*_0_ through *a* do not transfer exactly to other parameter directions.

#### 3.5.1 Acute phase infection elimination

The dynamics of infection progression in this study reveal how the interplay between viral replication, immune response, and lipid metabolism governs the transition between acute and chronic HCV infection. In order to study infection elimination by the immune response in acute HCV, we add memory cells *M* ^*∗*^ to the model:

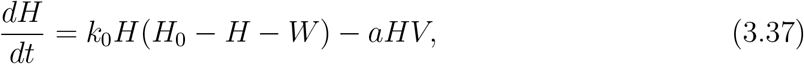

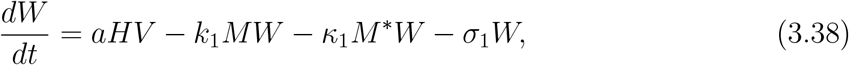

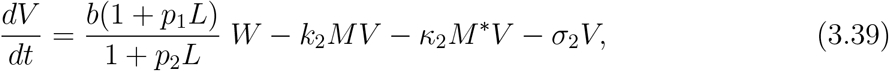

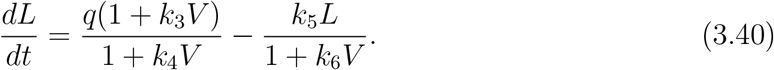

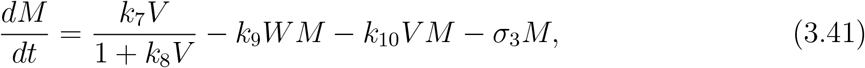

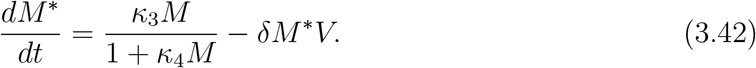

They differentiate from the memory cells *M* but they have a long life span. Therefore, we consider equation (3.42) without death term. These cells eliminate infected cells and neutralize/eliminate virus particle either directly or by intermediate of antibodies produced by B-lymphocytes.

In this formulation infected cells and viral particles are removed by both the transient immune factor *M* and the memory component *M* ^*∗*^. We then vary the memory depletion parameter *δ*. When *δ* = 0 and immune memory is not depleted by viral load or intrinsic decay, and the model exhibits acute infection followed by clearance. In this case the viral load reaches a transient peak and then decays to near zero.

For positive values of *δ*, immune memory is depleted during infection. A refined long-time sweep shows that the acute–chronic boundary is nonzero. Small positive values of *δ* still clear, but clearance becomes progressively slower as *δ* approaches the transition. On the grid considered, the last clearing value is *δ* = 2.40× 10^*−*7^ and the first chronic value is *δ* = 2.45× 10^*−*7^. Thus the transition should not be interpreted as occurring exactly at *δ* = 0; rather, a sufficiently large but still small memory-depletion rate is required to remove enough immune memory for persistence (Figure 8).

**Figure 8:**
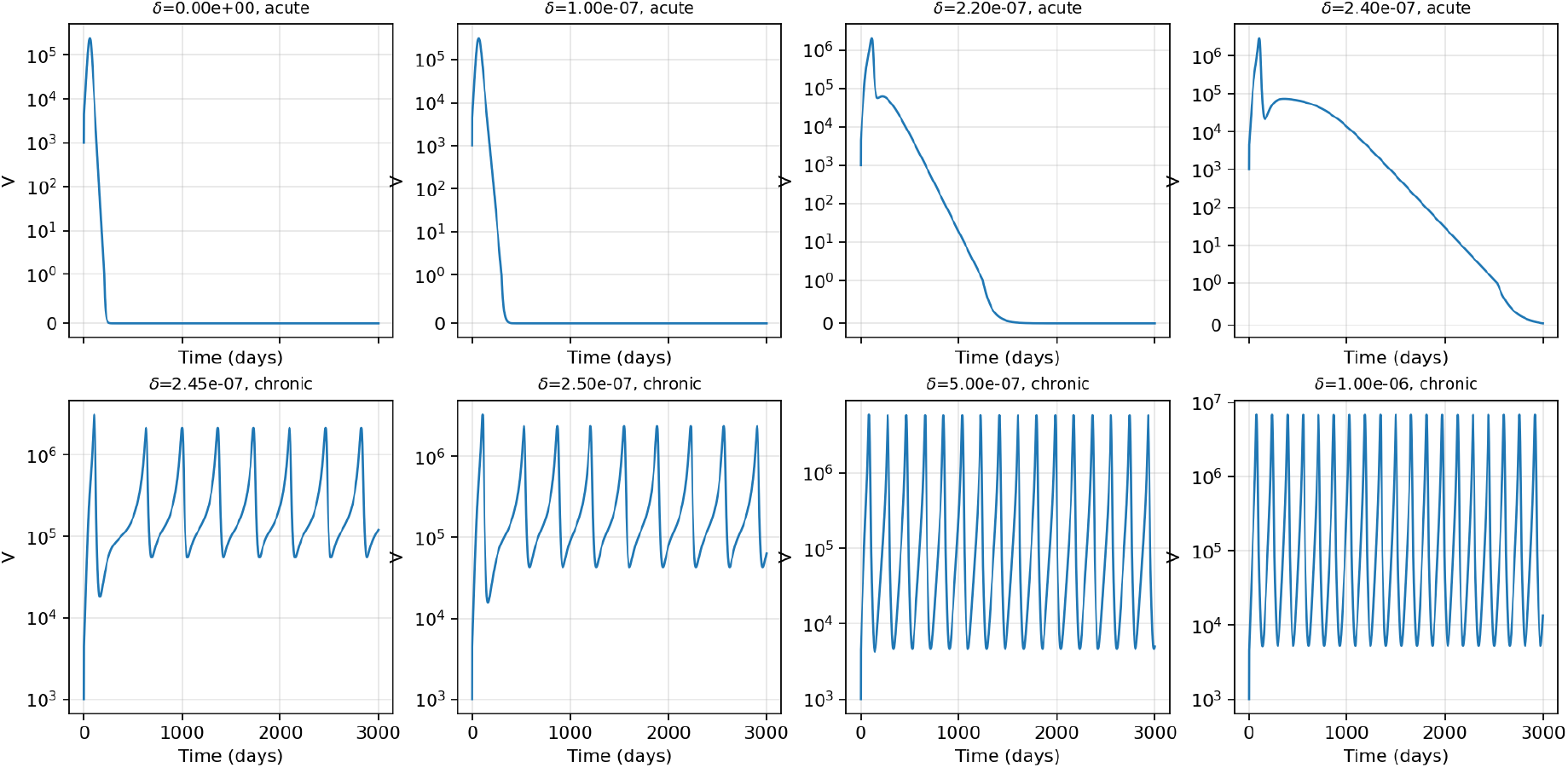
Viral-load trajectories for different values of the parameter *δ*, reflecting both memory cell depletion and elimination by the HCV virus. Infection clears at *δ* = 0 and for small positive values of *δ*, but the decay becomes slow near the boundary. Chronic persistent oscillations appear between *δ* = 2.40 × 10^*−*7^ and *δ* = 2.45 × 10^*−*7^ on the refined grid.

The acute–chronic transition is summarized in Figure 9. As *δ* increases, the peak viral load increases and the solution approaches a recurrent chronic regime with large viral excursions. The boundary band marks the interval between the last clearing sample and the first chronic sample, and the leading-eigenvalue panel shows the same change from negative to positive active stability margin. After the boundary is crossed, the dynamics settle into sustained recurrent oscillations with large peaks and nonzero viral tails. Thus, loss of immune memory can convert an acute resolving infection into chronic persistent infection, but only after a nonzero depletion threshold has been crossed.

**Figure 9:**
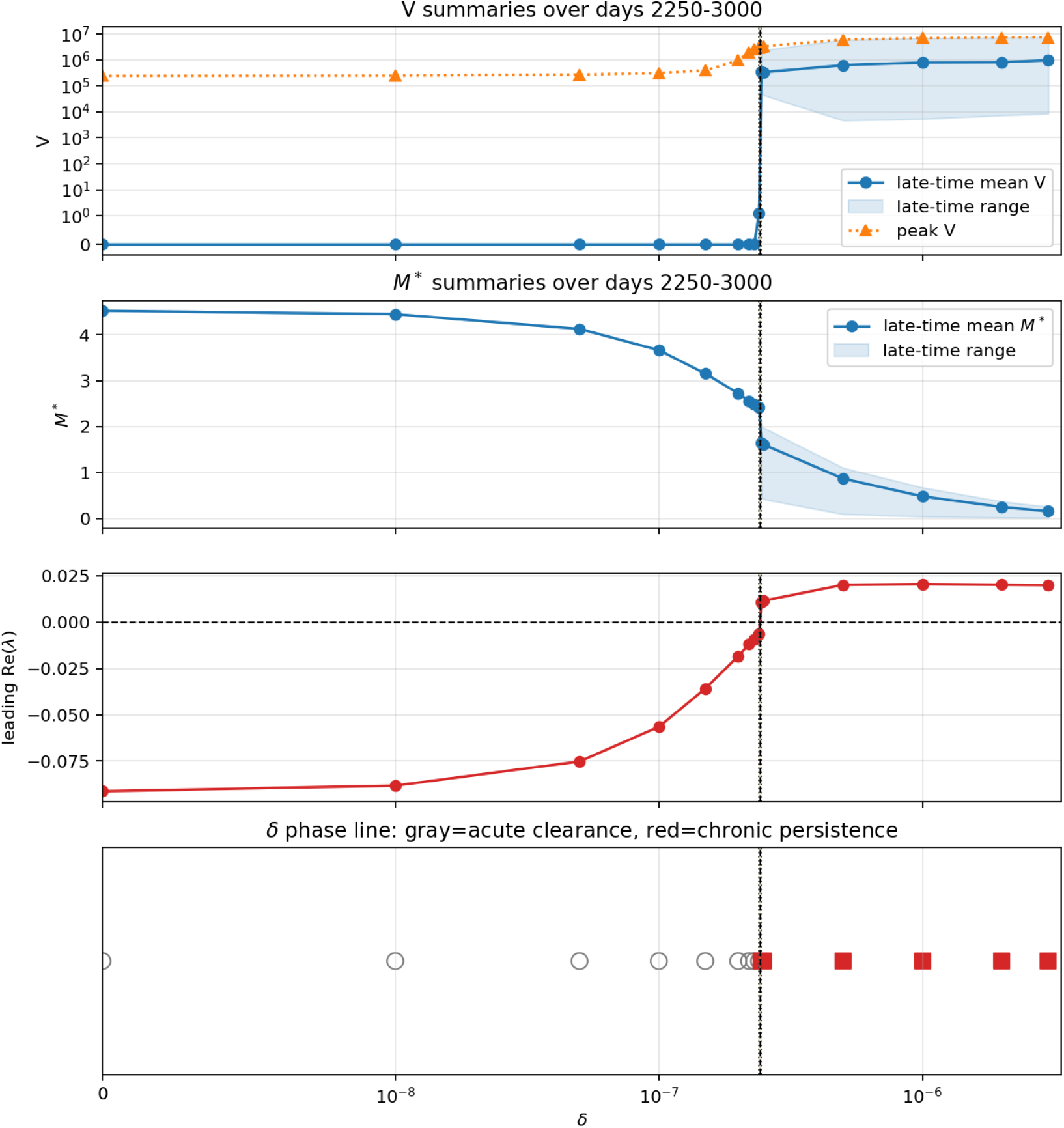
Summary of the refined *δ* sweep. The upper panel shows the tail mean, final value, and peak value of the viral load over a long-time integration. The second panel shows the residual immune memory *M* ^*∗*^, the third panel shows the leading active real eigenvalue, and the lower panel classifies the outcome as acute clearance or chronic persistence. The transition band lies between *δ* = 2.40 × 10^*−*7^ and *δ* = 2.45 × 10^*−*7^ on the grid considered.

#### 3.5.2 Sensitivity to immune killing and viral neutralization

We now vary the ratio

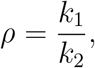

keeping *k*_2_ fixed. This ratio measures the relative strength of the *M* -mediated removal of infected cells compared with the *M* -mediated removal of free virus; the memory-mediated removal terms involving *κ*_1_ and *κ*_2_ are kept fixed. The results show that the dependence on

*ρ* is not monotone. For *ρ* = 10^3^ and *ρ* = 10^4^, the infection clears after a large transient peak. At *ρ* = 10^5^, the viral tail becomes large and the model enters a chronic oscillatory regime. At still larger ratio, *ρ* = 10^6^, the infection remains persistent but both the viral peak and tail are reduced compared with *ρ* = 10^5^ (Figure 10). These results suggest that changing the balance between the *M* -mediated removal pathways can qualitatively alter infection outcome, and that immune efficacy cannot be characterized by a single removal parameter alone.

**Figure 10:**
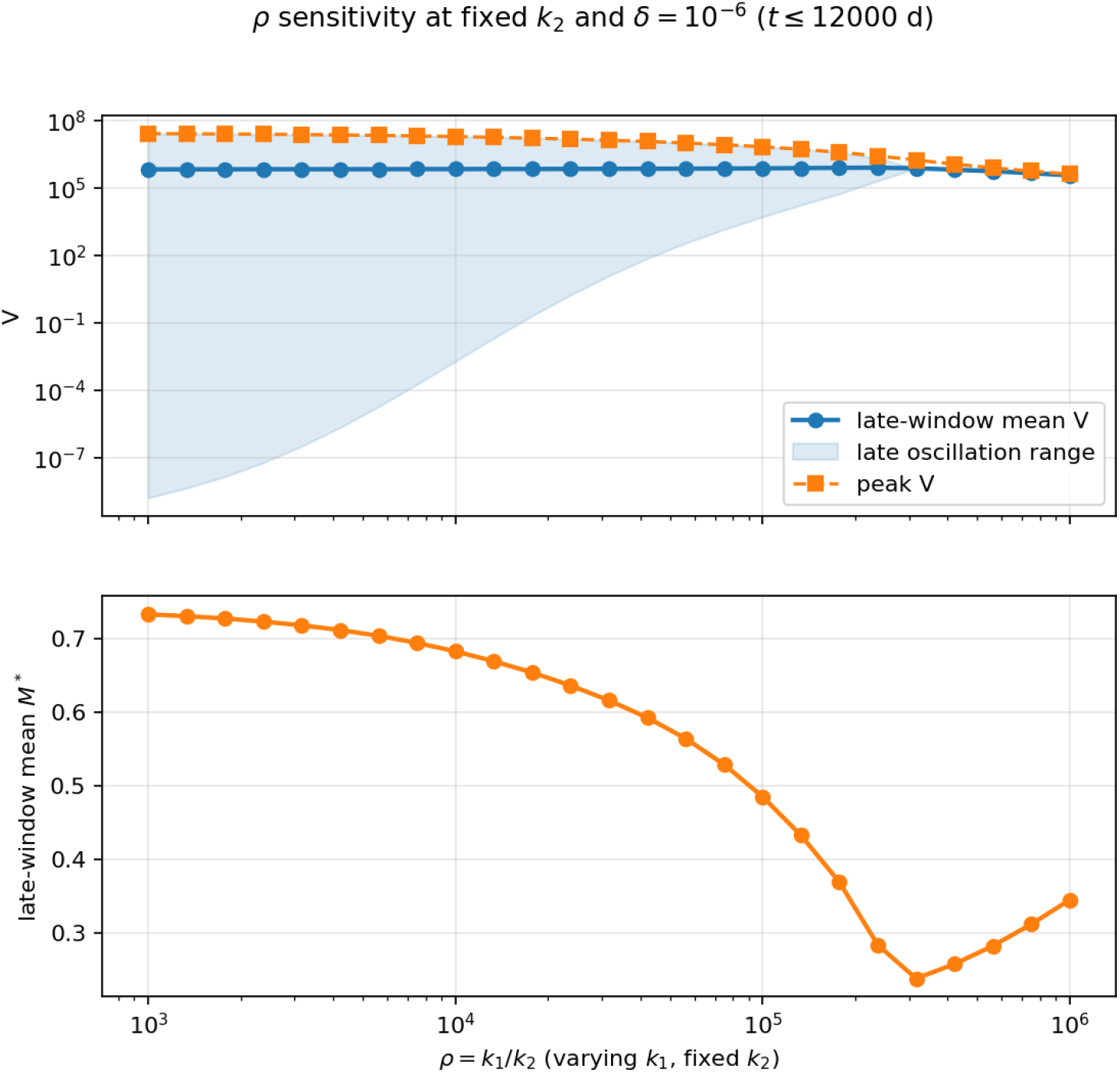
Sensitivity to the immune ratio *ρ* = *k*_1_*/k*_2_ at fixed *k*_2_ and *δ* = 10^*−*6^. The viral tail is small for *ρ* = 10^3^ and 10^4^, large for *ρ* = 10^5^, and reduced but persistent for *ρ* = 10^6^, showing that the relative balance of the *M* -mediated removal terms can change the infection outcome.

## 4 Dynamics for the PDE model

We now include the immune memory variable *M* ^*∗*^ in the system eq.(2.1)-eq.(2.5) to study infection elimination by the immune response in acute HCV and obtain the following PDE system.

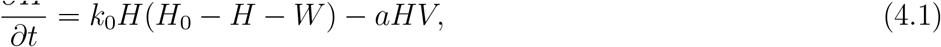

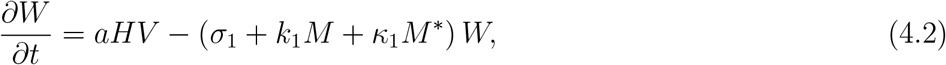

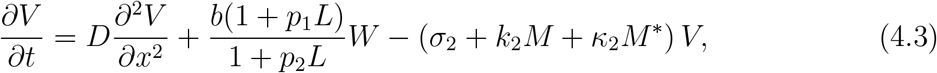

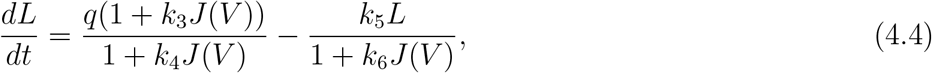

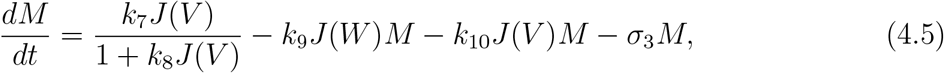

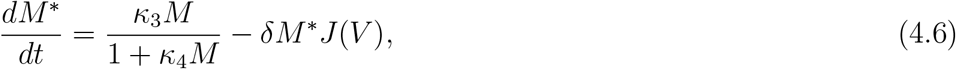

where, 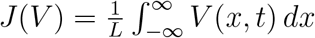,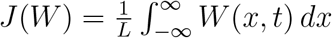, where *L* is the spatial length.

The numerical simulations for the PDE model eq.(4.1)-eq.(4.6) are performed on the one-dimensional domain Ω = [0, 1] subject to homogeneous Neumann boundary conditions. Unless otherwise stated, the parameter values are taken from the memory reference values in Table 3 and the other parameter values in the Table 4 and Table 5 corresponding to *R*_0_ = 1.94785. The initial conditions are *H*(*x*, 0) = *H*_0_ = 10^7^, *W* (*x*, 0) = 0, *L*(0) = 1, *M* (0) = 0, *M* ^*∗*^(0) = 0, and the initial virus distribution is given by 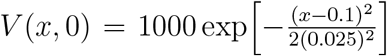 .

**Table 1:**
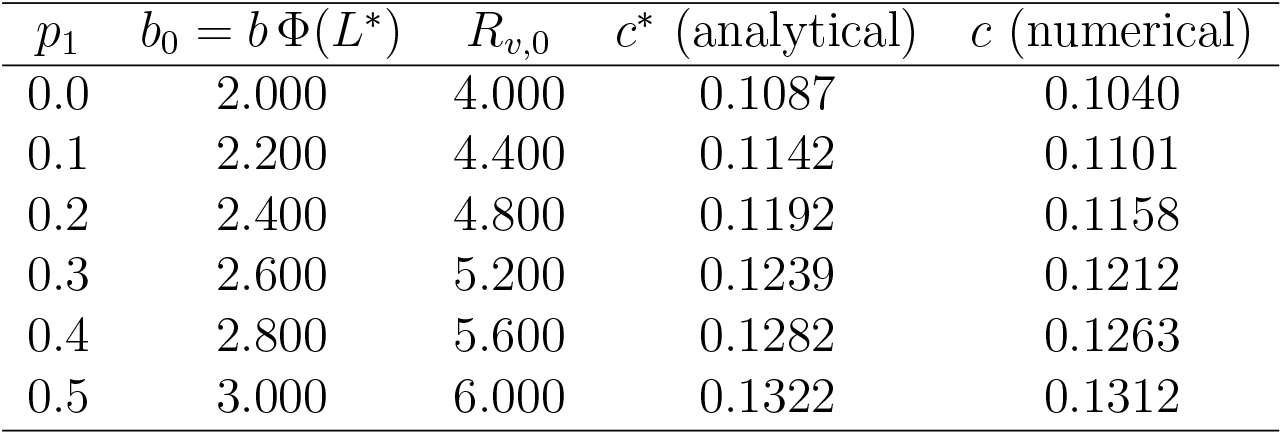
Values plotted in Figure 14.

| $p_1$ | $b_0 = b \Phi(L^*)$ | $R_{v,0}$ | $c^*$ (analytical) | $c$ (numerical) |
| --- | --- | --- | --- | --- |
| 0.0 | 2.000 | 4.000 | 0.1087 | 0.1040 |
| 0.1 | 2.200 | 4.400 | 0.1142 | 0.1101 |
| 0.2 | 2.400 | 4.800 | 0.1192 | 0.1158 |
| 0.3 | 2.600 | 5.200 | 0.1239 | 0.1212 |
| 0.4 | 2.800 | 5.600 | 0.1282 | 0.1263 |
| 0.5 | 3.000 | 6.000 | 0.1322 | 0.1312 |

**Table 2:** Literature-scale anchors and default numerical values used to set the main biological scale of the ODE experiments. Parameters without a listed anchor are retained as model coefficients rather than fitted literature quantities.

| Parameter | Meaning | Anchor range | Default value |
| --- | --- | --- | --- |
| $H_0$ | hepatocyte carrying scale | $4.6 \times 10^6$–$1.3 \times 10^7$ | $1.0 \times 10^7$ |
| $\sigma_1$ | infected-cell loss | 0.01–0.41 day<sup>−1</sup> | 0.1 |
| $\sigma_2$ | viral clearance | 5.5–8.3 day<sup>−1</sup> | 6.65 |
| $b$ | viral production | 0.1–45 | 2.12 |
| $p_1$ | lipid amplification | – | 0.3 |
| $p_2$ | lipid saturation | – | 0 |
| $q$ | lipid source scale | – | 0.5 |
| $k_5$ | lipid removal scale | – | 0.5 |

**Table 3:**
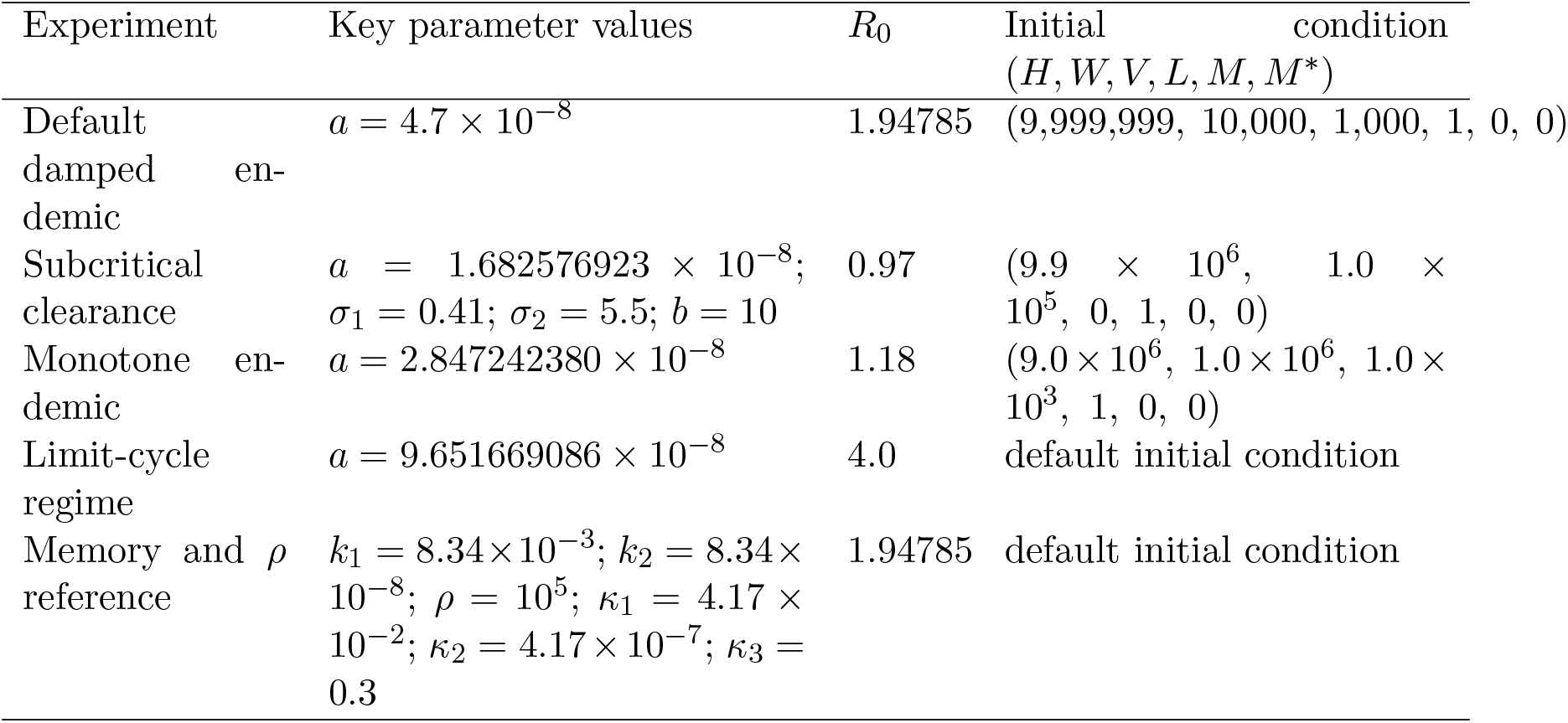
Parameter sets and initial conditions used for the main ODE experiments in the Results section. The first four rows correspond to the panels in Figure 2. Entries not shown are equal to the default values in Table 2 and the baseline configuration.

**Table 4:**
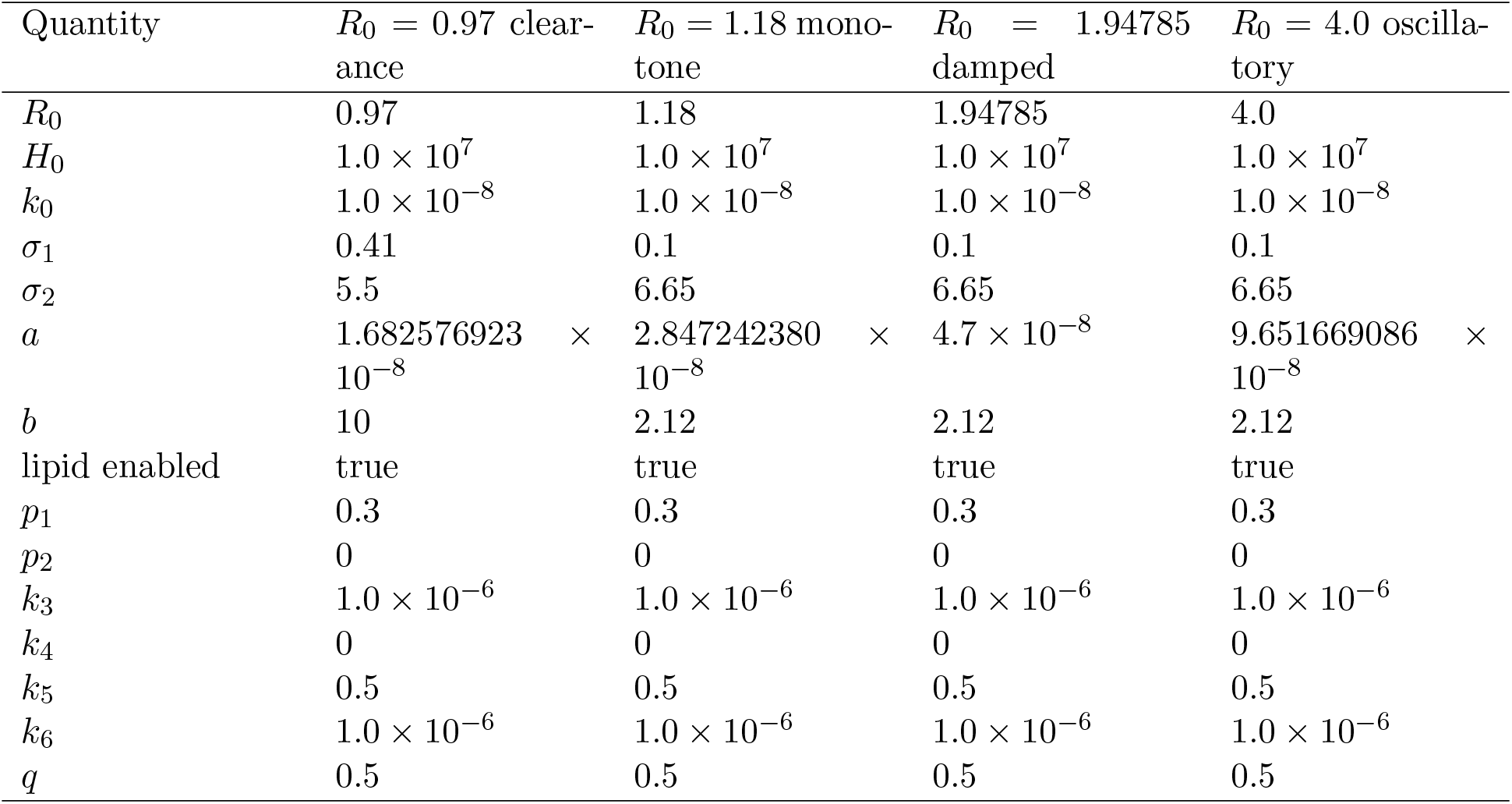
Core and lipid parameter values for the four ODE trajectory-catalog cases in Figure 2.

| Quantity | $R_0 = 0.97$ clear-<br>ance | $R_0 = 1.18$ mono-<br>tone | $R_0 = 1.94785$<br>damped | $R_0 = 4.0$ oscilla-<br>tory |
| --- | --- | --- | --- | --- |
| $R_0$ | 0.97 | 1.18 | 1.94785 | 4.0 |
| $H_0$ | $1.0 \times 10^7$ | $1.0 \times 10^7$ | $1.0 \times 10^7$ | $1.0 \times 10^7$ |
| $k_0$ | $1.0 \times 10^{-8}$ | $1.0 \times 10^{-8}$ | $1.0 \times 10^{-8}$ | $1.0 \times 10^{-8}$ |
| $\sigma_1$ | 0.41 | 0.1 | 0.1 | 0.1 |
| $\sigma_2$ | 5.5 | 6.65 | 6.65 | 6.65 |
| $a$ | $1.682576923 \times 10^{-8}$ | $2.847242380 \times 10^{-8}$ | $4.7 \times 10^{-8}$ | $9.651669086 \times 10^{-8}$ |
| $b$ | 10 | 2.12 | 2.12 | 2.12 |
| lipid enabled | true | true | true | true |
| $p_1$ | 0.3 | 0.3 | 0.3 | 0.3 |
| $p_2$ | 0 | 0 | 0 | 0 |
| $k_3$ | $1.0 \times 10^{-6}$ | $1.0 \times 10^{-6}$ | $1.0 \times 10^{-6}$ | $1.0 \times 10^{-6}$ |
| $k_4$ | 0 | 0 | 0 | 0 |
| $k_5$ | 0.5 | 0.5 | 0.5 | 0.5 |
| $k_6$ | $1.0 \times 10^{-6}$ | $1.0 \times 10^{-6}$ | $1.0 \times 10^{-6}$ | $1.0 \times 10^{-6}$ |
| $q$ | 0.5 | 0.5 | 0.5 | 0.5 |

**Table 5:**
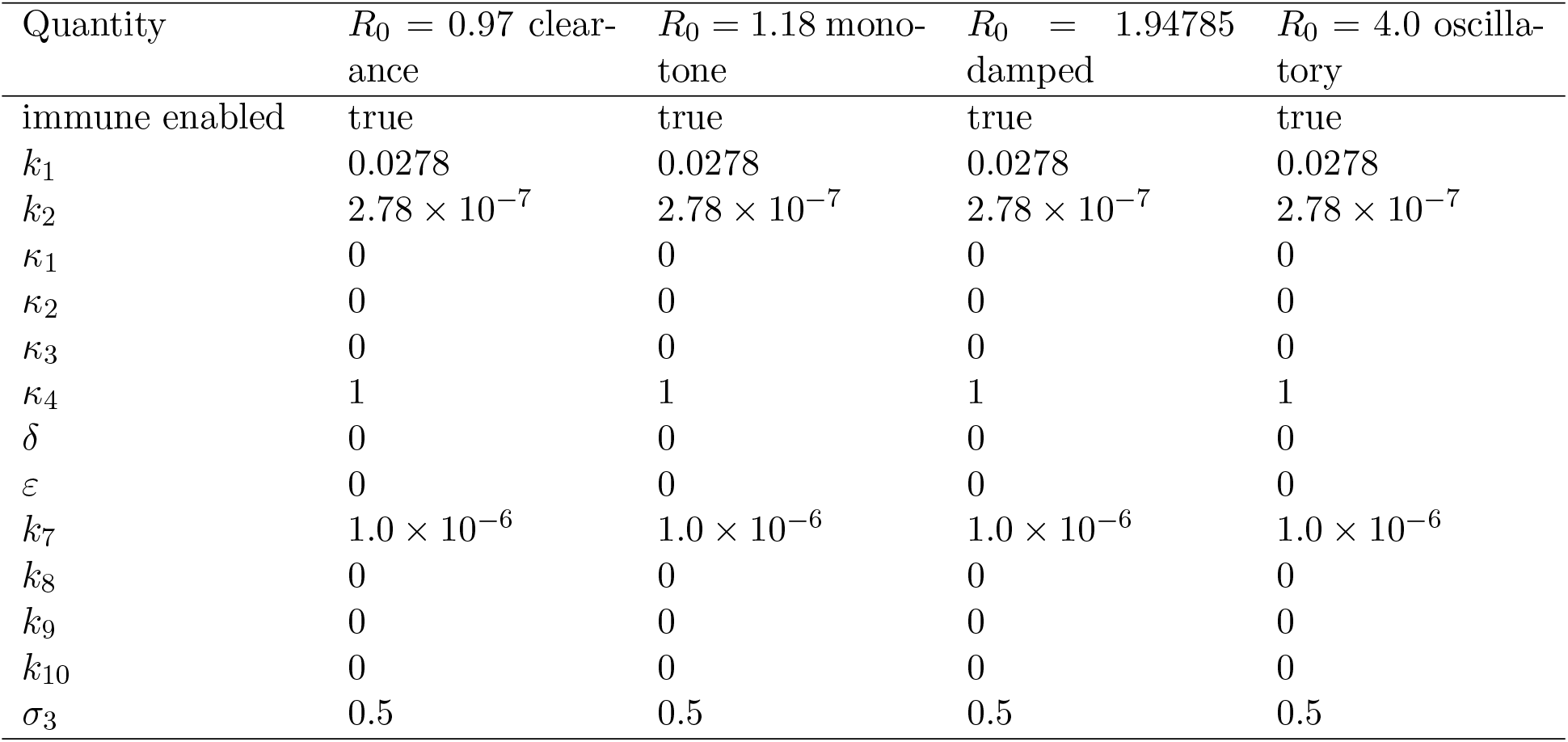
Immune parameter values for the four ODE trajectory-catalog cases in Figure 2.

| Quantity | $R_0 = 0.97$ clear-<br>ance | $R_0 = 1.18$ mono-<br>tone | $R_0 = 1.94785$<br>damped | $R_0 = 4.0$ oscilla-<br>tory |
| --- | --- | --- | --- | --- |
| immune enabled | true | true | true | true |
| $k_1$ | 0.0278 | 0.0278 | 0.0278 | 0.0278 |
| $k_2$ | $2.78 \times 10^{-7}$ | $2.78 \times 10^{-7}$ | $2.78 \times 10^{-7}$ | $2.78 \times 10^{-7}$ |
| $\kappa_1$ | 0 | 0 | 0 | 0 |
| $\kappa_2$ | 0 | 0 | 0 | 0 |
| $\kappa_3$ | 0 | 0 | 0 | 0 |
| $\kappa_4$ | 1 | 1 | 1 | 1 |
| $\delta$ | 0 | 0 | 0 | 0 |
| $\varepsilon$ | 0 | 0 | 0 | 0 |
| $k_7$ | $1.0 \times 10^{-6}$ | $1.0 \times 10^{-6}$ | $1.0 \times 10^{-6}$ | $1.0 \times 10^{-6}$ |
| $k_8$ | 0 | 0 | 0 | 0 |
| $k_9$ | 0 | 0 | 0 | 0 |
| $k_{10}$ | 0 | 0 | 0 | 0 |
| $\sigma_3$ | 0.5 | 0.5 | 0.5 | 0.5 |

**Table 6:**
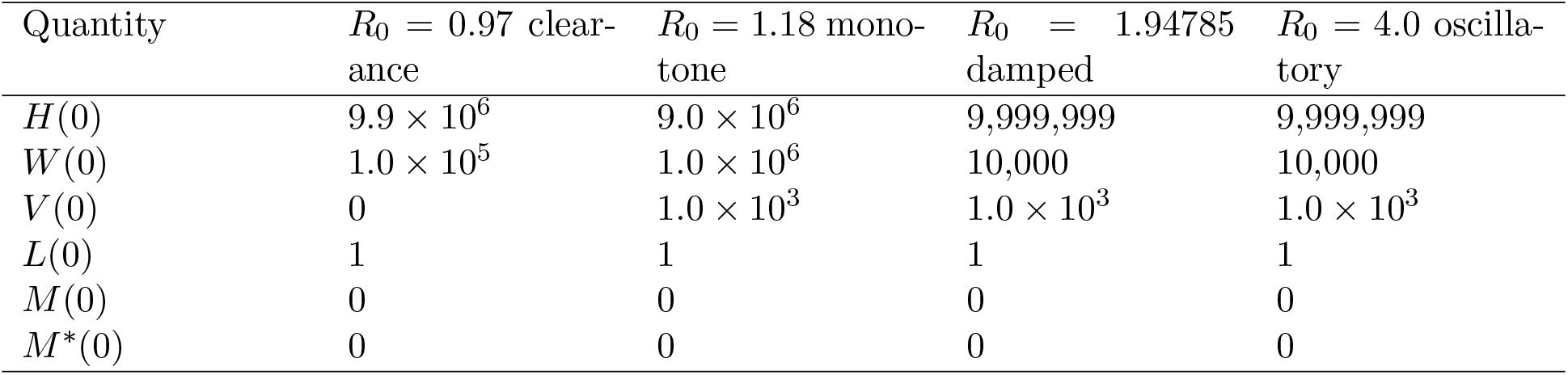
Initial conditions for the four ODE trajectory-catalog cases in Figure 2.

**Table 7:** Sweep directions and special parameter variations used in the Results figures.

| Study | Values or range | Interpretation |
| --- | --- | --- |
| Viral production $b$ | {0.3, 1.0, 2.12, 5.0, 15.0} | literature-bounded production sweep |
| Infected-cell loss $\sigma_1$ | {0.02, 0.08, 0.1, 0.2, 0.35} | literature-bounded loss sweep |
| Lipid amplification $p_1$ | {0.1, 0.2, 0.3, 0.4, 0.5} | lipid feedback sweep |
| Viral clearance $\sigma_2$ | 4.0–14.0 day<sup>−1</sup> | continuation diagnostic; partly extrapolative |
| Memory depletion $\delta$ | boundary bracket $2.40 \times 10^{-7}$–<br>$2.45 \times 10^{-7}$ | acute–chronic transition |
| Memory regularization $\varepsilon$ | 0 or $10^{-6}$ | native versus regularized memory equation |
| Immune ratio $\rho = k_1/k_2$ | {$10^3$, $10^4$, $10^5$, $10^6$} | $k_1$ varied at fixed $k_2$ |
| PDE seeding | spatial viral seed amplitude 1.0 | localized, shifted, step, and Gaussian profiles |

The results corresponding to the PDE model are shown in Figure 11 to Figure 13.

**Figure 11:**
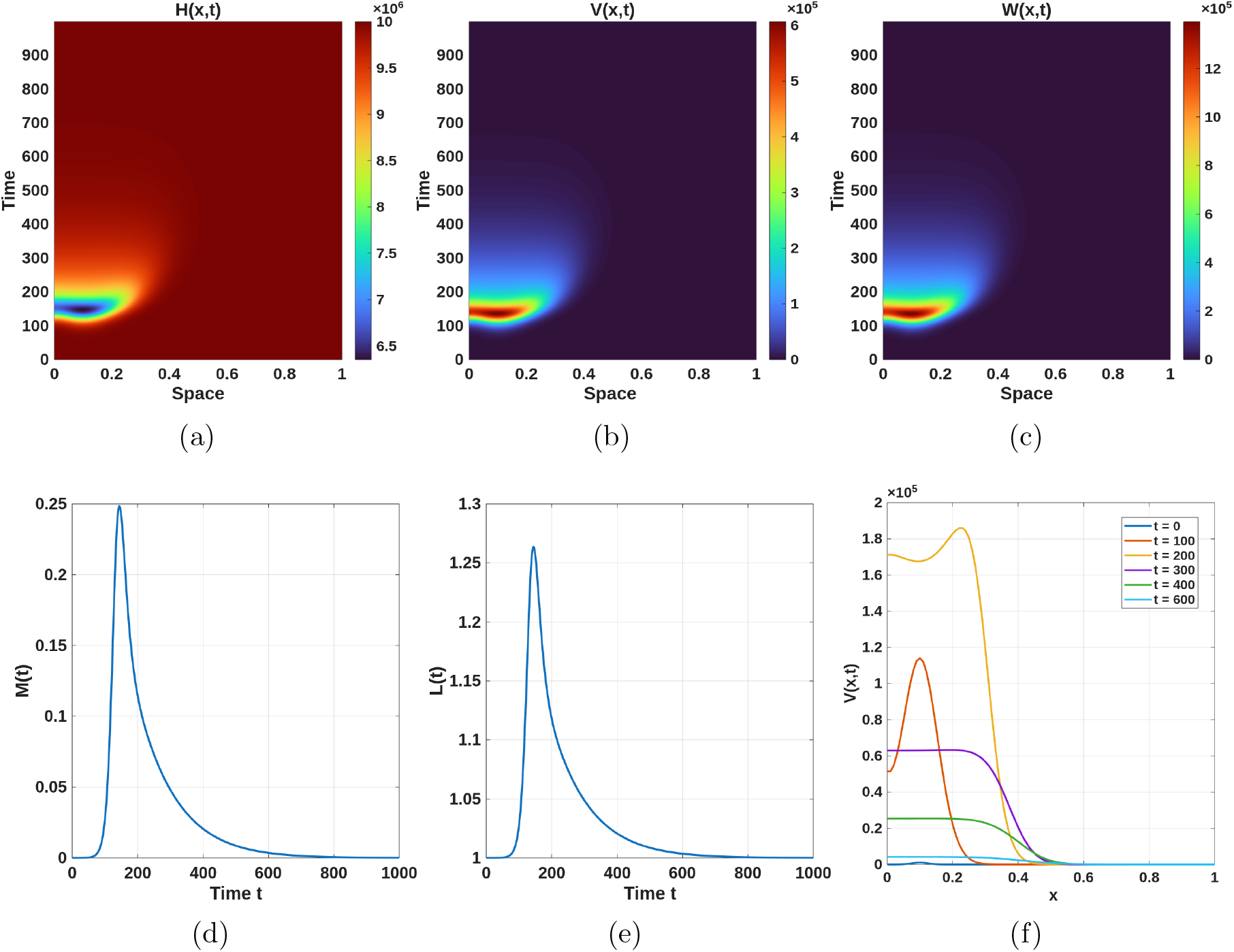
Panels (a)–(c) show the spatiotemporal evolution of *H*(*x, t*), *V* (*x, t*), and *W* (*x, t*), respectively. Panels (d) and (e) depict the temporal evolution of the nonlocal immune variables *M* (*t*) and *L*(*t*), while panel (f) shows the spatial profiles of the virus concentration *V* (*x, t*) at selected time instants. The parameter values, the initial conditions, and boundary conditions are mentioned in details in the main text.

### Burst and extinction scenario

Figure 11(a)–(c) show that the virus propagates in both space and time, reaching its peak around *t* ≈150 at the spatial location *x* ≈0.1. Moreover, the infection spreads in the neighbourhood of this peak, and after some time the infection dies out. It is evident that the spatial regions with a higher density of virus particles and infected cells correspond to regions with a lower density of healthy hepatocytes. However, as time progresses, both the virus and infected cell populations diminish, while the healthy cell population gradually recovers toward its disease-free level throughout the spatial domain. Panels (d) and (e) depict the temporal evolution of the nonlocal immune variables *M* (*t*) and *L*(*t*). Both variables exhibit a transient increase in response to the initial viral infection, attain a peak during the early stage of infection, and subsequently decline to their baseline levels as the viral load decreases. Panel (f) presents the spatial profiles of the virus concentration at selected time instants.

### Burst, propagation, and temporal oscillation scenario

If we increase the virus production rate, then the burst and elimination scenario shifts to a burst, propagation, and temporal oscillation scenario. From Figure 12, which corresponds to *b* = 2.4, it is observed that, due to the increase in *b*, the infection propagates and spreads throughout the entire spatial domain, eventually leading to sustained temporal oscillations. The temporal oscillation emerges once the spatial propagation has stopped. Moreover, a higher value of *b* results in a more rapid spread of infection across the spatial domain, and consequently, a larger number of temporal outbreaks are observed within the same time window (see Figures 12(b), (c)).

**Figure 12:**
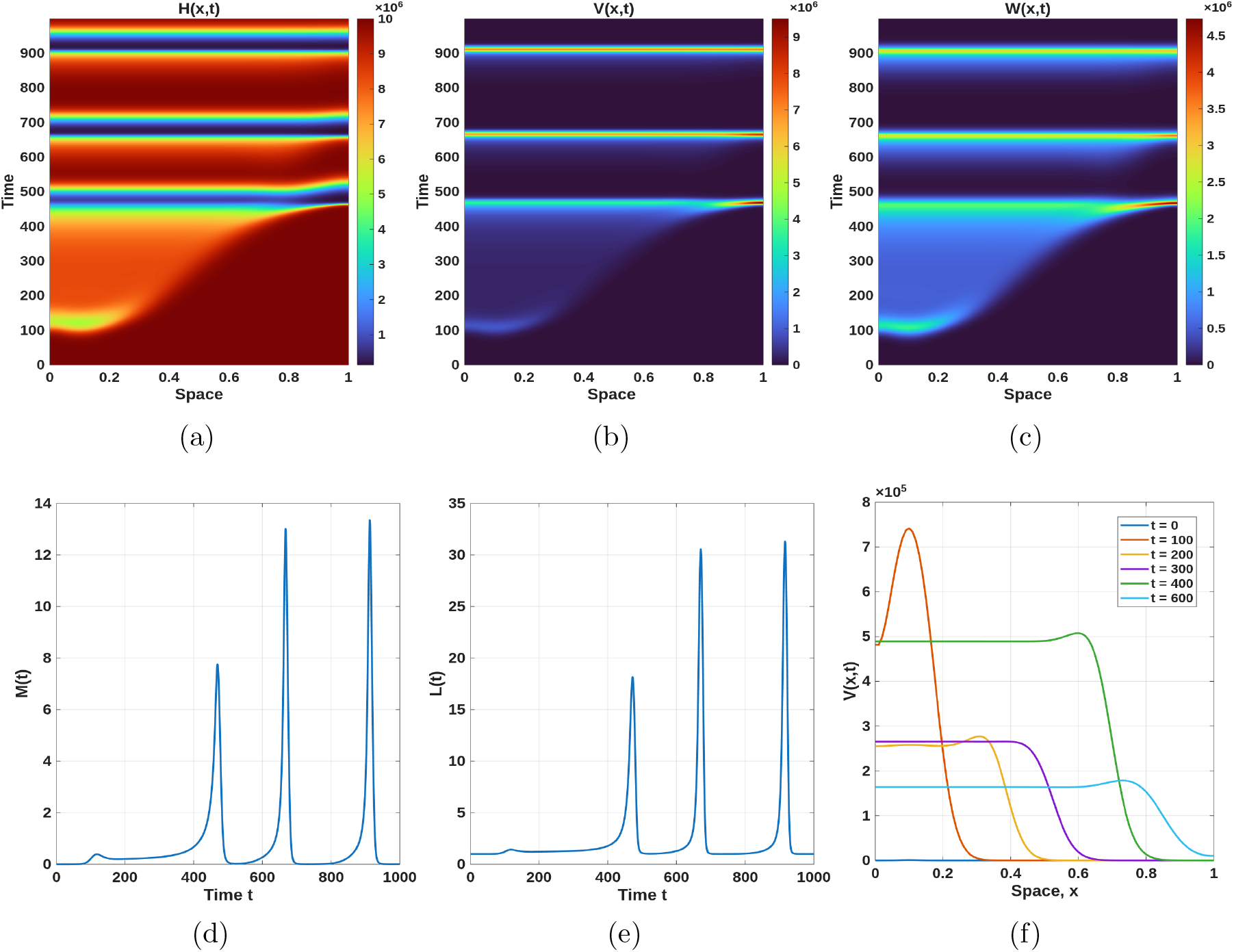
Panels (a)–(c) show the spatiotemporal evolution of *H*(*x, t*), *V* (*x, t*), and *W* (*x, t*), respectively. Panels (d) and (e) depict the temporal evolution of the nonlocal immune variables *M* (*t*) and *L*(*t*), while panel (f) shows the spatial profiles of the virus concentration *V* (*x, t*) at selected time instants. The value of *b* is taken as *b* = 2.4 and the the rest of the parameter values, the initial conditions, and boundary conditions are same as in Figure 11.

**Figure 13:**
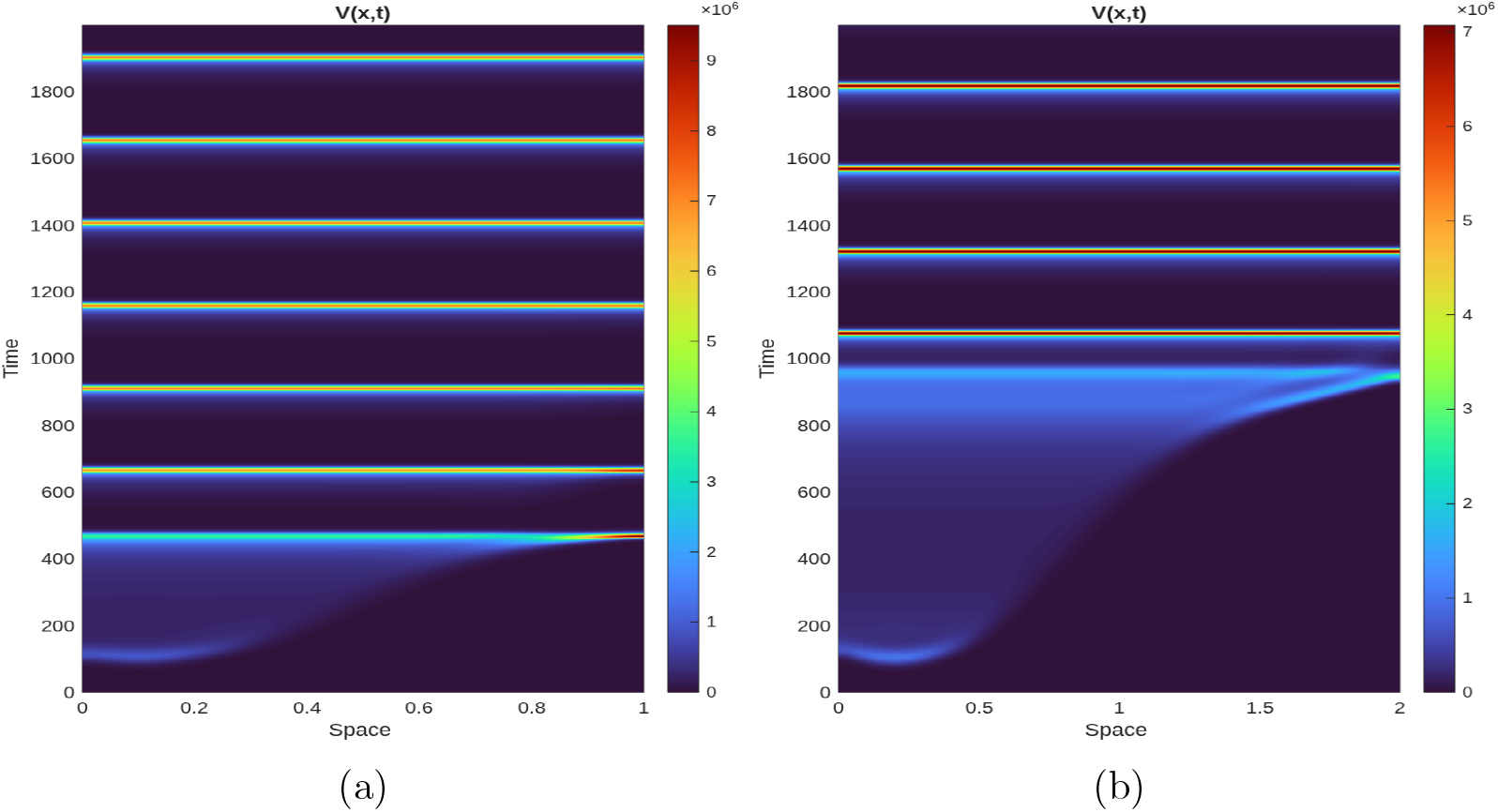
Plot of *V* (*x, t*) for different spatial lengths. Figure 13(a) and Figure 13(b) correspond to spatial length *x* = 1 and *x* = 2 respectively. All other parameter values and initial and boundary conditions as same as in Figure 12.

Moreover, since the temporal oscillations appear only after the infection has spread throughout the entire spatial domain, we also investigate the effect of the spatial domain length. In Figure 13, the virus compartment *V* (*x, t*) is plotted for the same parameter set as in Figure 12, but with different spatial domain lengths. Figures 13(a) and 13(b) correspond to spatial domain lengths of 1 and 2, respectively. It is observed that, under the same parameter setup, increasing the spatial domain length reduces the number of temporal outbreaks observed within a fixed time interval.

### 4.1 Analysis of infection spreading

Because lipids are produced and consumed locally within each hepatocyte, we keep the lipid balance in its local (per-cell) form, i.e. the form used in the ODE model, equation (3.4), rather than the spatially-averaged form of equation (2.4). Diffusion of lipids between cells is neglected, consistent with the rest of the model. With cell recovery neglected (*k*_0_ = 0) but the lipid-dependent viral production term retained, the spatial system analogous to (4.7)–(4.10) becomes

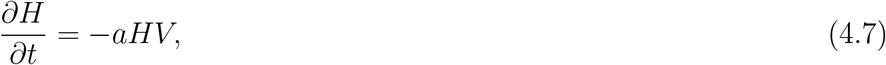

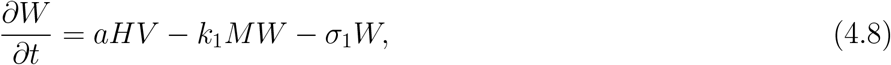

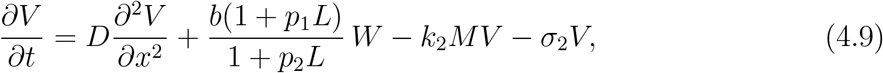

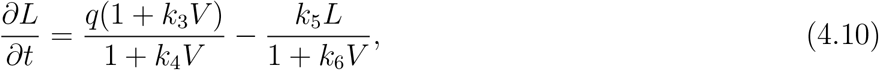

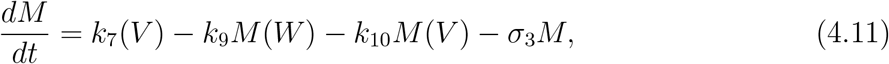

We look for a traveling-wave solution *H*(*x, t*) = *h*(*ξ*), *W* (*x, t*) = *w*(*ξ*), *V* (*x, t*) = *v*(*ξ*), *L*(*x, t*) = *l*(*ξ*), with *ξ* = *x* + *ct*, and *M* approaching a constant value *M* consistent with the wave. We obtain the following traveling-wave system

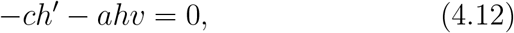

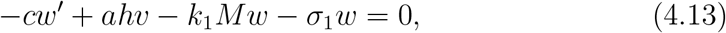

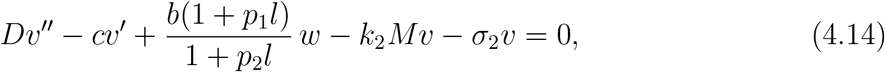

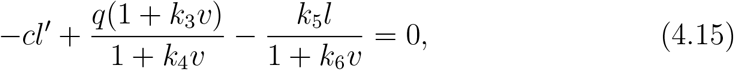

completed by the same algebraic balance for *M* as in the main text,

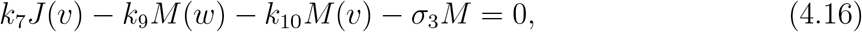

and the limits at infinity

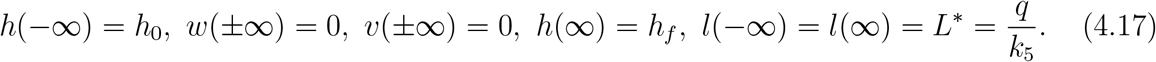

The condition *l*(±∞ ) = *L*^*∗*^ follows directly from (4.15): since *v*(±∞ ) = 0 and *l* approaches a constant at both ends (*l*^*′*^→ 0), the limiting balance 0 = *q* −*k*_5_*l* forces *l* →*q/k*_5_ on both sides of the wave. Thus *l*(*ξ*), like *v*(*ξ*) and *w*(*ξ*), is a localized perturbation of a background level, here the disease-free lipid equilibrium *L*^*∗*^ already identified in the Appendix.

### 4.2 Quasi-steady-state closure for the lipid profile

Equation (4.15) couples *l*(*ξ*) to *v*(*ξ*) through a first-order, non-diffusive relation. When intracellular lipid turnover is fast compared with the time the front takes to sweep a fixed point in space (i.e. *c l*^*′*^ is small relative to the two terms on the right of (4.15)), *l* equilibrates quasi-instantaneously to the local viral load,

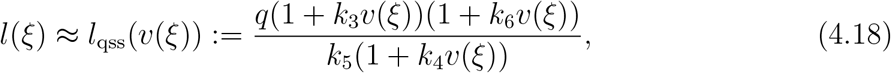

which is exactly the stationary-point formula (3.12) evaluated pointwise along the wave. This closure reduces the system to three coupled equations for (*h, w, v*), with the lipid feedback entering (4.14) through the explicit function Φ(*l*_qss_(*v*)), where

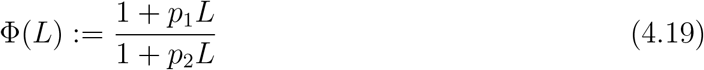

is the lipid amplification factor already used in the Appendix (there evaluated at *L*^*∗*^). Note Φ(*L*^*∗*^) is precisely Φ(*l*_qss_(0)), i.e. the value of the amplification factor at the disease-free lipid level.

#### Lemma 2.

*If k*_3_ + *k*_6_ ≥ *k*_4_ *(in particular if k*_3_ ≥ *k*_4_, *as assumed for the model), then l*_qss_(*v*) *is nondecreasing in v for v* ≥ 0, *with l*_qss_(0) = *L*^*∗*^. *Consequently, if in addition p*_1_ ≥ *p*_2_, *then* Φ(*l*_qss_(*v*)) ≥ Φ(*L*^*∗*^) *for all v* ≥ 0.

*Proof*. Write 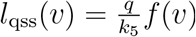 with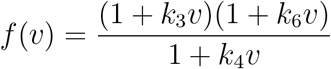. A direct computation gives

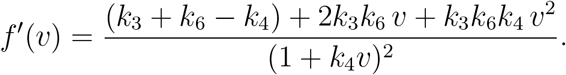

Since *k*_3_, *k*_4_, *k*_6_ ≥ 0, the terms 2*k*_3_*k*_6_*v* and *k*_3_*k*_6_*k*_4_*v*^2^ are nonnegative for *v* ≥ 0; hence *f* ^*′*^(*v*) ≥ 0 for all *v* ≥ 0 whenever *k*_3_ + *k*_6_ ≥ *k*_4_. This proves the first claim. For the second, Φ^*′*^(*L*) = (*p*_1_ − *p*_2_)*/*(1 + *p*_2_*L*)^2^ ≥ 0 when *p*_1_ ≥ *p*_2_, so Φ is nondecreasing; composing with the nondecreasing map *v* ↦ *l*_qss_(*v*) ≥ *L*^*∗*^ gives Φ(*l*_qss_(*v*)) ≥ Φ(*L*^*∗*^).

#### 4.2.1 Values of *h*_*f*_ and viral load with lipid feedback

Equations (4.12)–(4.13) do not involve *L* directly, so the first two integral relations of the main text carry over unchanged. Dividing (4.12) by *h* and integrating,

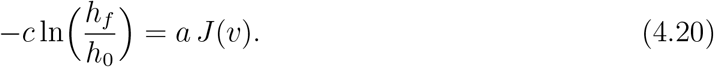

Summing (4.12) and (4.13) and integrating, using *w*(±∞) = 0,

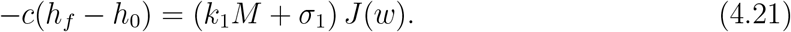

Integrating (4.14) over the whole line (the diffusive and advective terms vanish because *v, v*^*′*^ → 0 at both ends) gives, in place of (4.18),

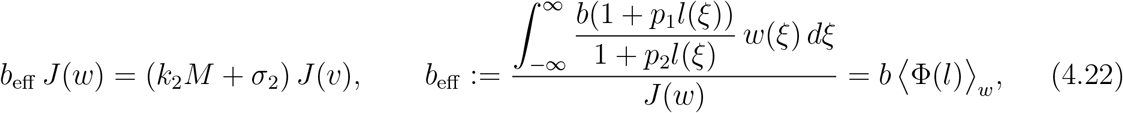

i.e. the constant *b* of the lipid-free analysis is replaced by its *w*-weighted average along the wave, *b*⟨Φ(*l*)⟩_*w*_. By Lemma 2 (under the QSS closure (4.18)), *l*(*ξ*) ≥ *L*^*∗*^ everywhere, so

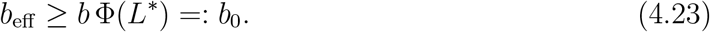

As in the main text, set *k*_9_ = *k*_10_ = 0 (immune factors are not depleted by their interaction with infected cells or virus). Then (4.16) gives *M* = (*k*_7_*/σ*_3_)*J* (*v*), and (4.22) gives

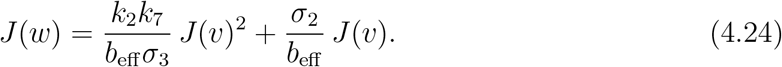

Substituting *M* and (4.24) into (4.21), with *h*_*f*_ − *h*_0_ = *h*_0_(*η* − 1) and *η* := *h*_*f*_ */h*_0_,

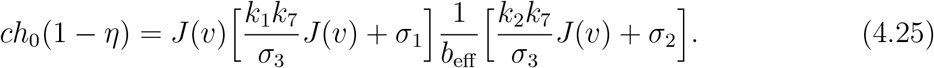

Using (4.20) to write *J* (*v*) = − (*c/a*) ln *η* and dividing by *c* (*c* ≠ 0), exactly as in the derivation of (4.20),

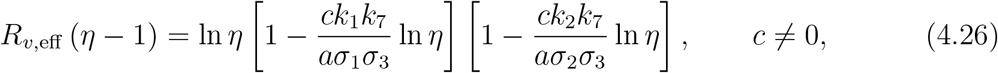

where

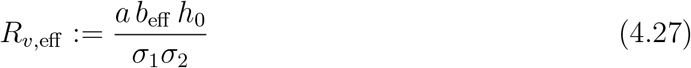

is an *effective* viral replication number that now depends on the lipid feedback through *b*_eff_ . Equation (4.26) has exactly the form of (4.20), so the same argument as the Proposition in the main text applies verbatim:

##### Proposition 3.

*Equation* (4.26) *has a solution η* ∈ (0, 1) *if and only if R*_*v*,eff_ *>* 1.

Because *R*_*v*,eff_ is not known in closed form (it depends on the full nonlinear wave profile through *b*_eff_ ), the bound (4.23) yields an explicit, fully computable *sufficient* condition for the front to exist:

##### Corollary 3.

*Let R*_*v*,0_ := *ab*_0_*h*_0_*/*(*σ*_1_*σ*_2_) = *ab* Φ(*L*^*∗*^)*h*_0_*/*(*σ*_1_*σ*_2_), *with* Φ(*L*^*∗*^) = (1 + *p*_1_*L*^*∗*^)*/*(1 + *p*_2_*L*^*∗*^) *and L*^*∗*^ = *q/k*_5_. *If R*_*v*,0_ *>* 1, *i*.*e*.

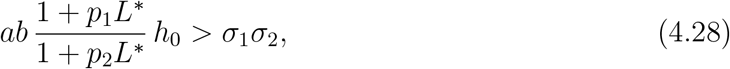

*then R*_*v*,eff_ ≥ *R*_*v*,0_ *>* 1 *and equation* (4.26) *has a solution η* ∈ (0, 1).

Since Φ(*L*^*∗*^) ≥1 whenever *p*_1_ ≥*p*_2_ and *L*^*∗*^ *>* 0, Corollary 3 shows that a nonzero baseline (pre-infection) hepatic lipid level *L*^*∗*^ already lowers the threshold for spatial invasion.

### 4.3 Minimal wave speed with lipid feedback

The minimal wave speed is governed by the linearization about the leading edge of the front, where *H* ≈*h*_0_ and, since *v* ≈ 0 ahead of the front, *L* ≈ *L*^*∗*^ by (4.18). Linearizing (4.13)–(4.14) about this state (i.e. *h* = *h*_0_, *l* = *L*^*∗*^) gives

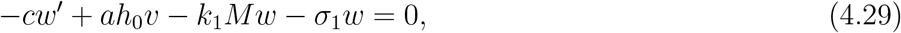

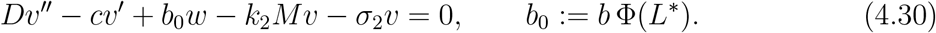

These are identical in structure to (4.22)–(4.23) with *b* replaced by *b*_0_: the leading edge only “feels” the lipid amplification factor evaluated at the disease-free level *L*^*∗*^, because to linear order in *v* the deviation of *l* from *L*^*∗*^ is itself of higher order. Seeking *w* = *p*_1_*e*^*−λξ*^, *v* = *p*_2_*e*^*−λξ*^ and setting *µ* = *λc* as in the main text, the same elimination of *p*_1_, *p*_2_ yields the dispersion relation

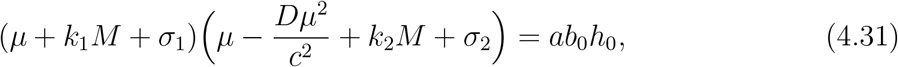

so that

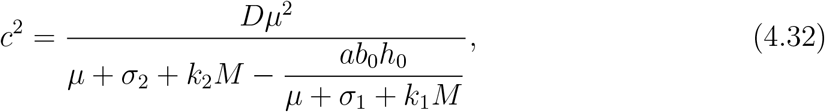

and the minimal wave speed is

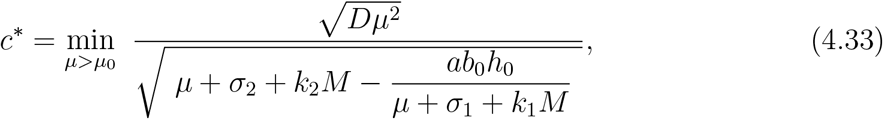

where *µ*_0_ *>* 0 is the value at which the denominator vanishes; this reduces exactly to formula (4.24) of the main text when *L*^*∗*^ = 0 (i.e. *b*_0_ = *b*). A minimum exists, exactly as in condition (4.25), provided

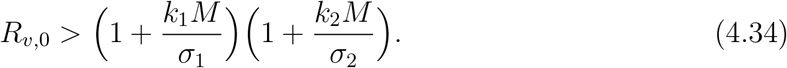

Summarizing, viral infection spreads in the tissue if *R >* 1 + *P*, with 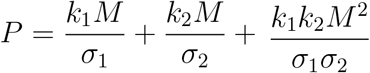 with *R*_*v*,0_ = *ab* Φ(*L*^*∗*^)*h*_0_*/*(*σ*_1_*σ*_2_).

### 4.4 Numerical validation of the wave speed

We compare the analytically obtained wave speed, formula (4.33), with the wave speed computed from direct numerical simulation of the full nonlinear system (4.7)–(4.10). Following the “first approximation” recommended in the main text, we set *k*_1_ = *k*_2_ = 0 (no immune response), so that *M* drops out of (4.33) and the comparison isolates the effect of the lipid feedback alone, through *b*_0_ = *b* Φ(*L*^*∗*^). The remaining parameters are *a* = 0.5, *σ*_1_ = *σ*_2_ = 0.5, *D* = 0.01, *b* = 2, *h*_0_ = 1, *p*_2_ = 0, *q* = *k*_5_ = 1 (so *L*^*∗*^ = 1), *k*_3_ = 1, *k*_4_ = 0.2, *k*_6_ = 0, and the lipid amplification parameter *p*_1_ is varied over [0, 0.5].

The PDE system is solved with a semi-implicit finite-difference scheme (explicit diffusion, implicit linear removal terms) on a domain of length 900 with no-flux boundaries, starting from a localized viral seed on an otherwise uninfected reservoir, *H*(*x*, 0) = *h*_0_. The numerical wave speed is estimated from the constant-speed drift of the location of the viral peak, arg max_*x*_ *V* (*x, t*), over the interval *t* ∈ [400, 800], after the initial transient has died out.

Table 1 lists the underlying values. The two curves increase together as *p*_1_ grows, confirming that the lipid-corrected formula (4.33) correctly tracks how baseline lipid amplification speeds up spatial invasion, even though the numerical simulation retains the full nonlinear feedback *b*(1 + *p*_1_*L*)*/*(1 + *p*_2_*L*) (with *L* evolving *above L*^*∗*^ behind the front, per Lemma 2) rather than the frozen value *b*_0_ used in the linearization. The small residual gap between the two curves is consistent with the well-known slow (algebraic/logarithmic) relaxation of a pulled front toward its asymptotic linear-spreading speed, rather than with any structural disagreement between (4.33) and the simulated dynamics.

## 5 Data fitting to HCV RNA trajectories

We fitted the ODE viral state *V* (*t*) to HCV RNA trajectories by estimating a subset of model parameters together with an observation-scale parameter. The data source is [8]. The model output was converted to the fitted measurement scale by

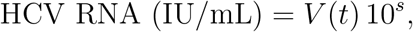

where *s* is the fitted log-scale conversion. The fitted observable was

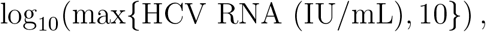

with 10 IU/mL used as the detection-floor value.

For persistent infection, the preferred fit varied the core viral parameters *a, b, σ*_1_, and *σ*_2_, the initial viral load *V* (0), the observation-scale parameter *s*, and a constrained infected-cell fraction *f*_inf_ . The initial infected-cell and uninfected-cell populations were then set by

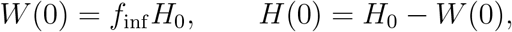

which enforces *H*(0) + *W* (0) = *H*_0_. This fit gave RMSE = 0.502 on the log_10_ HCV RNA scale and *R*_0_ = 9.33.

The fitted persistent-infection parameter values were

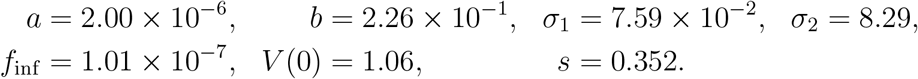

Thus *H*(0) = 9.61 × 10^6^ and W(0) = 3.94 × 10^5^.

The persistent-infection fit gives *R*_0_ = 9.33 together with an extremely small fitted infected-cell fraction, *f*_inf_ = *W* (0)*/H*_0_ ≈ 1.01 × 10^*−*7^. Both features have direct physiological interpretations: A minute initial infected-cell fraction is consistent with the biology of HCV transmission, in which a real exposure event (needle-stick, transfusion, perinatal transmission) seeds only a tiny inoculum relative to the full hepatocyte pool, not a macroscopic fraction of the liver. A large fitted *R*_0_ ≫ 1 reflects strong intrinsic viral replicative fitness, consistent with the clinical observation that a majority of acute HCV infections progress to chronicity rather than resolving spontaneously.

The fitted trajectory itself is not a simple monotone rise to a plateau: it shows an early partial decline (months 3–4) followed by a rebound (month 5) before settling into a fluctuating chronic level (Figure 15). This qualitative shape is the clinical signature of an early, partially effective innate/adaptive immune response that transiently suppresses the virus, followed by incomplete immune control or a lipid-feedback-driven replication rebound – precisely the mechanism responsible for the damped and sustained oscillatory regimes identified analytically in Section 3.3 via the destabilizing term *κ* = *ψ*^*′*^(*V* ^*∗*^)*W* ^*∗*^.

**Figure 14:**
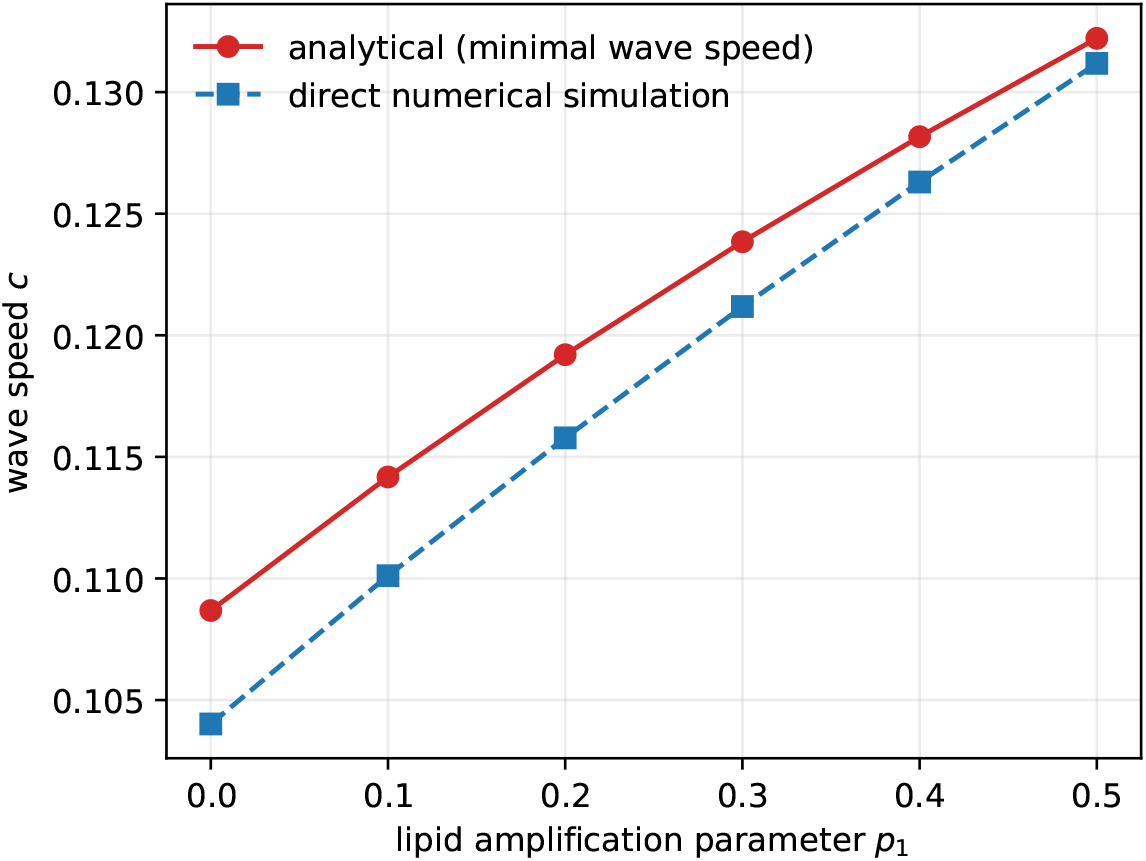
Comparison between the analytically obtained wave speed using formula (4.33) (red circles) and the wave speed obtained from direct numerical simulation of system (4.7)– (4.10) (blue squares), for different values of the lipid amplification parameter *p*_1_, with *k*_1_ = *k*_2_ = 0 and other parameter values as follows: *a* = 0.5, *σ*_1_ = *σ*_2_ = 0.5, *b* = 2, *h*_0_ = 1, *D* = 0.01, *q* = *k*_5_ = 1 (*L*^*∗*^ = 1), *k*_3_ = 1, *k*_4_ = 0.2, *k*_6_ = 0, *p*_2_ = 0. Both curves increase with *p*_1_ at comparable rates, and agree to within about 2–4% over the whole range, validating the leading-edge prediction that the minimal wave speed depends on the lipid feedback only through the disease-free amplification factor Φ(*L*^*∗*^) = 1 + *p*_1_*L*^*∗*^.

**Figure 15:**
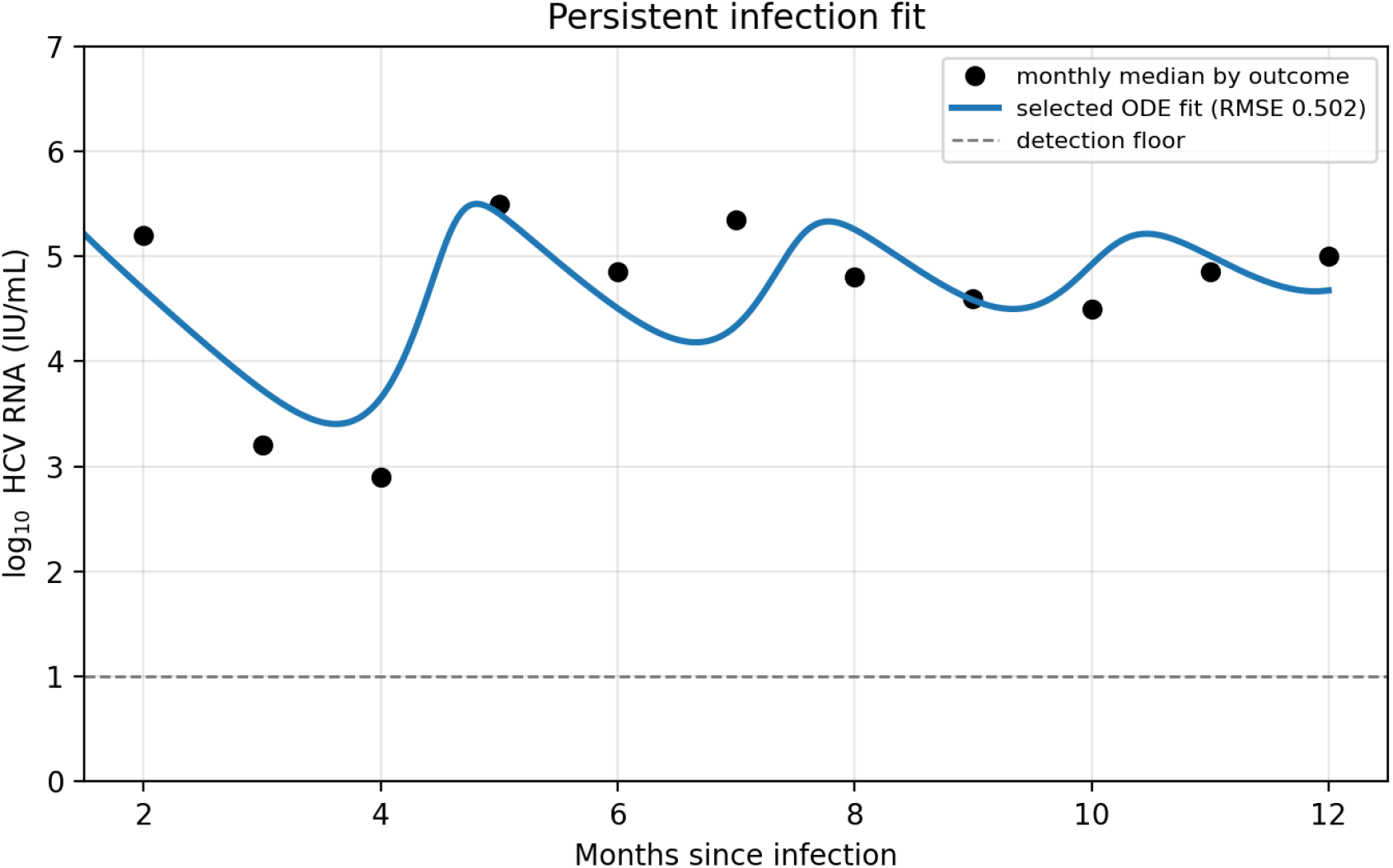
ODE fit to the persistent-infection HCV RNA trajectory. The fitted viral state *V* (*t*) is converted to log_10_ HCV RNA IU/mL using a fitted observation-scale parameter. The preferred fit varies core viral parameters, initial viral load, and a constrained infected-cell fraction satisfying *H*(0) + *W* (0) = *H*_0_.

For spontaneous clearance as in Figure (16), the fitting procedure varied *a, b, σ*_1_, *σ*_2_, *k*_1_, *k*_2_, *k*_7_, *W* (0), *V* (0), and the observation-scale parameter*s*. The unweighted clearance fit gave RMSE = 0.123 on the log10 HCV RNA scale.

The clearance fit achieves a substantially lower error (RMSE = 0.123 versus 0.502 for the persistent fit) and reproduces a clean, monotonic decline of viral RNA to the assay detection floor by approximately month 5. This matches the well-documented clinical pattern that spontaneous clearers typically resolve viremia within the first several months of acute infection, and corresponds to a parameter regime with effective *R*_0_ *<* 1, so that the disease-free equilibrium *E*_0_ is the attracting state and the trajectory decays toward it rather than toward an endemic branch 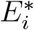.

## 6 Discussion and Conclusion

In this work, we develop a comprehensive mathematical framework that integrates hepatitis C virus (HCV) infection dynamics with intracellular lipid metabolism to investigate the mechanisms underlying viral-induced steatosis. By coupling viral kinetics with lipid regulatory pathways and immune responses, the model captures the bidirectional feedback between virus replication and host metabolic processes. Specifically, HCV-driven alterations in lipid influx were shown to promote intracellular lipid accumulation, while elevated lipid levels enhance viral replication, establishing a reinforcing feedback loop central to disease progression.

The results show that within the model, the intracellular lipid pool is not a passive marker riding along with the viral load at equilibrium, it is an active co-determinant of *whether* an endemic state can exist (via 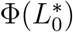 raising *R*_0_), *how large* it is (via the amplification of the effective production rate *b*Φ(*L*^*∗*^(*V* ))), and *whether it persists in a stable steady state or instead relapses* (via the destabilizing term *κ*). Physiologically this corresponds to the hypothesis that HCV-induced steatosis is not simply a histological consequence of chronic infection but an active participant in a metabolic–viral feedback loop that helps commit the liver to a chronic, and potentially fluctuating, infection state, providing a mechanistic, equilibrium-level explanation for the clinical association between the degree of steatosis and both viral load and disease severity.

The analytical and numerical investigations show that the outcome of infection is fundamentally governed by threshold quantities analogous to a viral replication number. When this threshold exceeds unity, infection persists and leads to a stable or oscillatory endemic state, whereas values below the threshold result in viral clearance. The framework further highlights the emergence of multiple dynamical regimes including transient infection, chronic persistence, damped oscillations, and sustained oscillatory behavior demonstrating the spectrum of possible disease trajectories within a unified model structure. These regimes are controlled by key biological parameters such as viral production, immune-mediated clearance, and lipid-mediated amplification, underscoring the interplay of virological, metabolic, and immunological factors.

A central insight of this study is the critical role of lipid feedback in shaping infection dynamics. Increases in lipid accumulation not only reflect metabolic dysregulation but actively promote viral replication, thereby amplifying infection severity and persistence. This mechanistic link provides a quantitative explanation for clinical observations associating steatosis with higher viral loads and more severe disease progression. The model thus supports the hypothesis that targeting lipid metabolic pathways such as lipogenesis may complement antiviral therapies by disrupting the positive feedback loop between lipids and viral replication. The incorporation of immune dynamics, including a simplified representation of immune response and memory effects, offers further insight into acute versus chronic infection outcomes. The results indicate that a nonzero threshold of immune memory depletion governs the transition from viral clearance to chronic persistence. This finding suggests that even subtle impairments in immune memory maintenance can substantially alter long-term infection outcomes, providing a plausible mechanistic explanation for variability observed in patient responses. Moreover, sensitivity to the relative strength of immune mediated removal pathways emphasizes that immune efficacy cannot be characterized by a single parameter but depends on the balance of multiple processes.

Spatial extensions of the model demonstrate that localized viral seeding evolves into tissue wide infection patterns through diffusion, while maintaining sensitivity to parameter values controlling viral production and immune response. These results highlight the importance of spatial heterogeneity in early infection dynamics, although long-term behavior is primarily determined by systemic interactions captured in the model.

The proposed framework is further supported by its ability to reproduce experimentally observed HCV RNA trajectories through data fitting, capturing both persistent infection and spontaneous clearance scenarios. This validation underscores the utility of the model as a predictive tool for interpreting clinical data and exploring therapeutic strategies.

Despite its strengths, the model incorporates several simplifying assumptions, including aggregated immune variables and reduced representations of lipid metabolism. Future work could extend this framework by incorporating more detailed immune cell populations, spatial heterogeneity in liver architecture, and genotype-specific viral effects. Additionally, integrating patient-specific data and pharmacological interventions would enhance its applicability to personalized medicine and treatment optimization. In conclusion, this study provides a comprehensive mathematical dynamics of HCV infection coupled with host lipid metabolism. By elucidating the feedback mechanisms linking viral replication, lipid accumulation, and immune response, the model advances our understanding of HCV-induced steatosis and identifies key determinants of infection outcome. Beyond HCV, the framework offers a generalizable approach for studying virus metabolism interactions in liver disease and may inform the development of combined antiviral and metabolic therapeutic strategies.

Future work on this framework include extension of the model to incorporate a more detailed immune response, explicitly distinguishing between innate and adaptive components such as cytotoxic T cells, B cells, and cytokine signaling networks, which would allow a more mechanistic study of immune-mediated viral clearance and persistence.

## 7 Appendix

### 7.1 Parameterization of the numerical experiments

The numerical experiments use one model structure but several deliberately chosen parameterizations. This separation is important because the figures in the Results section are not all generated from a single implicit default parameter set. The baseline configuration is used to display the damped endemic regime and to define the main viral, lipid, and immune scales. Additional configurations are then obtained either by tuning the infection rate *a* to a specified value of *R*_0_, by varying one parameter along a sweep direction, or by replacing the default immune coefficients with the reference immune-memory preset used in the acute–chronic transition study.

At the disease-free state, the lipid equilibrium is

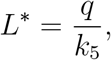

and the lipid-dependent production factor is

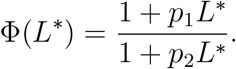

The replication number used to organize the ODE simulations is therefore

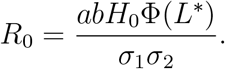

For the default numerical configuration, *L*^*∗*^ = 1, Φ(*L*^*∗*^) = 1.3, and *R*_0_ = 1.94785, with *k*_8_ = 0 in the immune stimulation term. This is the parameterization used for the damped oscillatory endemic case in Figure 2. The subcritical, monotone endemic, and large-*R*_0_ oscillatory examples are constructed from the same formula by changing *a*, except for the subcritical clearance case, which also uses the literature-scale values *σ*_1_ = 0.41, *σ*_2_ = 5.5, and *b* = 10.

The principal ODE parameter sets and initial conditions are summarized in Table 3. The first four rows are the four parameterizations used in the trajectory catalog in Figure 2. The state is *y* = (*H, W, V, L, M, M* ^*∗*^). The default initial condition contains a small infected cell population and a small viral inoculum, while the subcritical and monotone endemic examples use larger initial infected-cell populations to make the transient dynamics visible on the plotted time scale.

The *R*_0_ value reported for the memory and *ρ* reference row is the same zero-memory virus–lipid scale defined above. It is not an effective reproduction number that incorporates retained *M* ^*∗*^ at a memory-protected disease-free state.

For complete reproducibility of Figure 2, Tables 4–6 list every parameter used by the implemented ODE right-hand side, together with the initial condition for each of the four trajectory-catalog cases. Parameters that are stored but not used by the right-hand side are not included in these tables and are discussed below.

The parameter sweeps in Figures **??**–6 and 7 are designed to show how the model moves between qualitative regimes. The *b* and *σ*_1_ sweeps remain within the stated biological anchor ranges. The *σ*_2_ row in the threshold-map figure is different: it is a numerical continuation diagnostic over 4.0–14.0 day^*−*1^, so the upper part of the scan extends beyond the literature anchor 5.5–8.3 day^*−*1^. Consequently, the transitions near *σ*_2_≈10.9 and *σ*_2_ ≈ 12.9 should be interpreted as model-extrapolation thresholds for that parameter direction, not as literature-bounded biological estimates.

The corresponding phase portraits in the (*W, V* ) plane show how these regimes arise geometrically (Figure 17). In the clearance case, the trajectory makes a large excursion and then returns toward the disease-free state. In the stable endemic cases, trajectories approach a positive fixed point, either directly or through a spiral. In the unstable oscillatory case, the trajectory does not settle to the equilibrium but instead remains on a recurrent orbit. These projections are useful for interpreting both the ODE model and the spatial simulations, because the PDE system uses the same local reaction terms for infection and viral production, with diffusion added to the viral equation.

**Figure 16:**
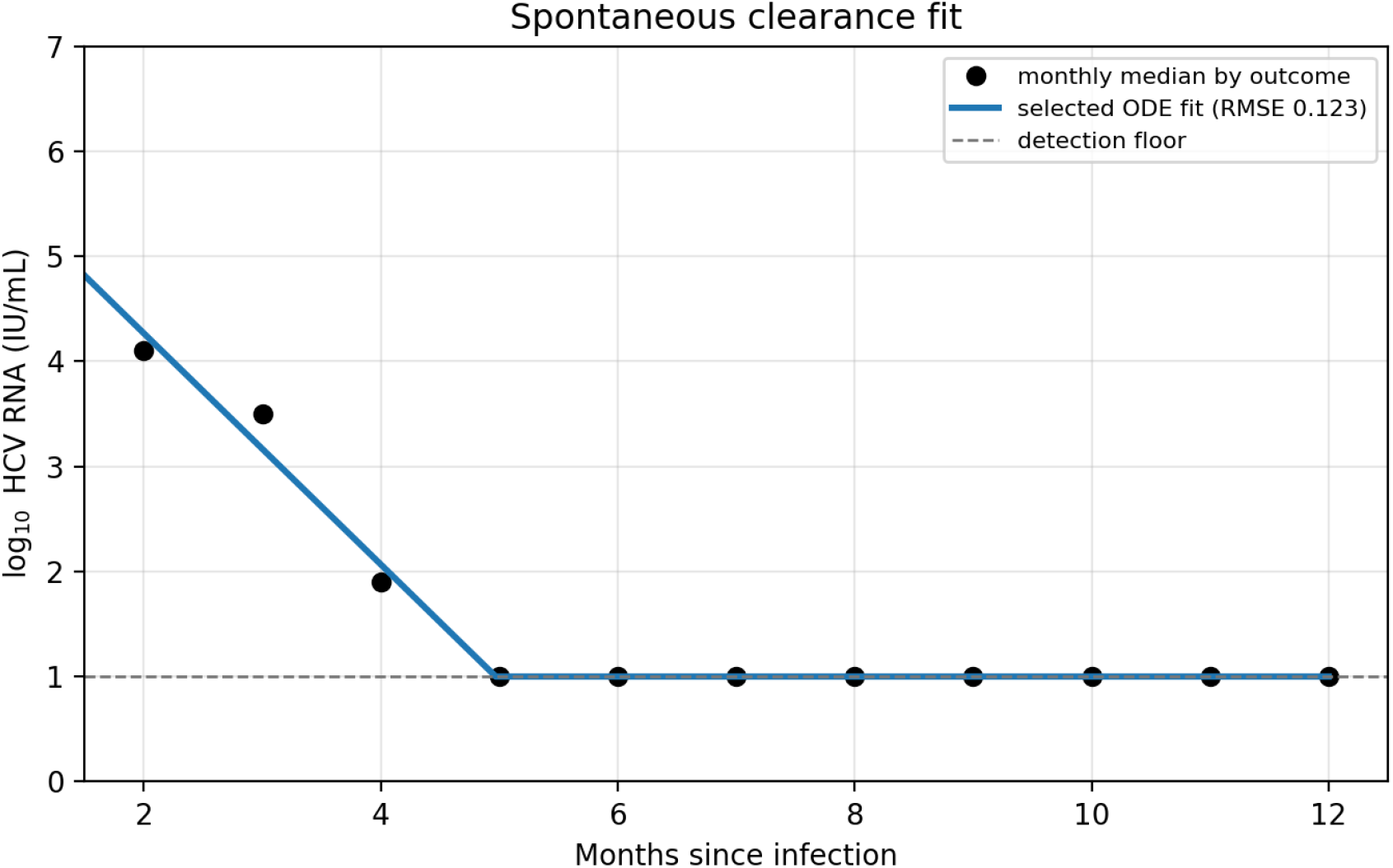
ODE fit to the spontaneous-clearance HCV RNA trajectory. The unweighted fit varies core viral, immune, initial-condition, and observation scale parameters and is evaluated on the log10 HCV RNA IU/mL scale.

**Figure 17:**
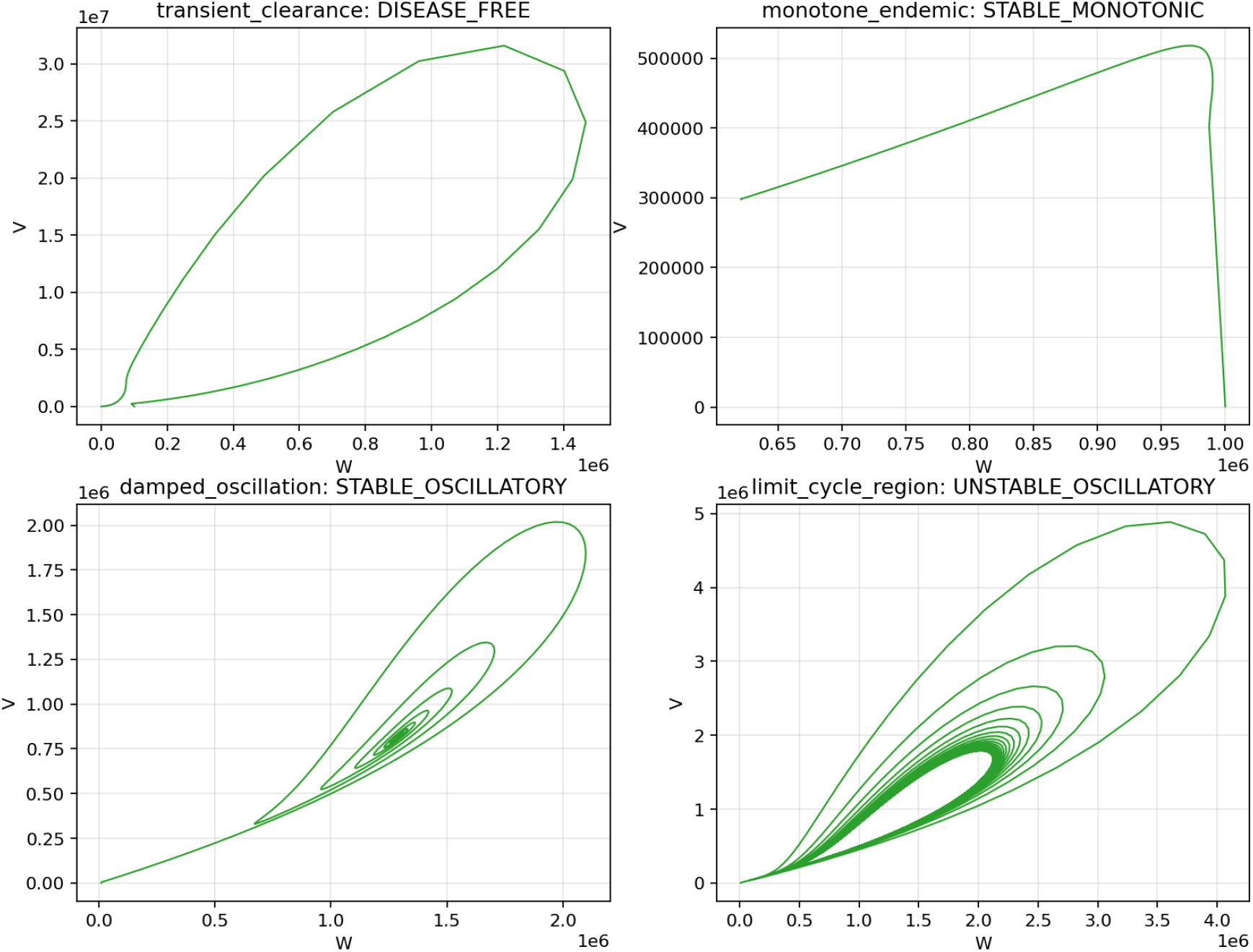
Phase portraits in the (*W, V* ) plane for the four representative ODE regimes. The plots show clearance as return toward the disease-free state, stable endemic infection as convergence to a positive equilibrium, damped oscillations as a spiral toward the endemic equilibrium, and persistent oscillations as recurrent motion around an unstable endemic equilibrium.

